# Fragmented sleep during pregnancy induces inflammation and tryptophan-kynurenine pathway metabolism: Importance of elevated fetal brain kynurenic acid

**DOI:** 10.64898/2026.06.24.734303

**Authors:** Courtney J. Wright, J. Hunter Cox, Snezana Milosavljevic, Homayoun Valafar, Norma Frizzell, Ana Pocivavsek

**Author notes:** Corresponding Author*: Ana Pocivavsek, PhD, Pharmacology, Physiology, and Neuroscience USC Floyd School of Medicine, Columbia, SC 29209, USA.

## Abstract

Maternal sleep disturbance is an underrecognized risk factor for adverse offspring outcomes. Prolonged sleep disruption can elicit inflammation, an established risk factor for neuropsychiatric disorders in offspring. Sleep disruptions and inflammation elevate tryptophan degradation via the kynurenine pathway (KP), increasing kynurenic acid (KYNA), a metabolite that inhibits glutamatergic and cholinergic neurotransmission and may thereby affect neurodevelopment. Because KYNA is elevated in the brains of individuals with neurodevelopmental psychotic illnesses, we hypothesize that prenatal KYNA elevation may represent a mechanism link between disturbed maternal sleep, inflammation, and adverse offspring neurodevelopmental health. To test this hypothesis, we employed a novel maternal sleep fragmentation (SleepFrag) paradigm during the final week of gestation. We found that six days of SleepFrag increased maternal plasma inflammatory markers, placental KP metabolism, sex-specific placental inflammation, and fetal brain KP metabolism, including elevated KYNA, without altering KP metabolism in maternal plasma or brain. A parallel embryonic kynurenine (EKyn) model was tested to increase prenatal KP metabolism via a maternal kynurenine-supplemented diet. EKyn increased maternal plasma kynurenine and KYNA, and fetal brain KYNA, with a male-specific increase in fetal brain KYNA-to-kynurenine ratio, despite minimal effects on maternal sleep-wake architecture or inflammation. Together, these findings identify elevated fetal brain KYNA as a convergent outcome through which maternal sleep disruption, inflammation, and KP activation may influence sex-specific neurodevelopment. They further support the EKyn model as a translational tool for isolating consequences of increased prenatal KP metabolism. Protecting maternal sleep and stabilizing fetal brain KYNA levels may promote long-term offspring brain health.

## 1. Introduction

Maternal inflammation during pregnancy is a well-established risk factor for impaired neurodevelopment and increased psychiatric disease risk in offspring (Estes and McAllister, 2016; Han et al., 2021). Although most often studied in the context of maternal infection, inflammation can also be induced by common physiological insults during pregnancy, including metabolic stressors, chronic stress, and sleep disruption (Hantsoo et al., 2019; Kwon et al., 2022; Méndez Leal et al., 2023; Okun, 2019; Wright et al., 2023b). Inadequate maternal sleep is a highly prevalent and significant concern during pregnancy. Clinical evidence increasingly links insufficient maternal sleep, particularly when combined with inflammation, with adverse health outcomes for pregnant individuals and their offspring (Facco et al., 2018; Huang et al., 2025; Lugo-Candelas et al., 2023; Nevarez-Brewster et al., 2024; Nevarez-Brewster et al., 2025; Okun, 2019; Yang et al., 2022; Zhang et al., 2025). Preclinical studies in rodents further show that experimental sleep deprivation increases pro-inflammatory cytokines and, when conducted during pregnancy, leads to persistent behavioral deficits in offspring, including sleep disturbances and cognitive dysfunction relevant to neurodevelopmental and psychiatric conditions (Irwin et al., 2016; Satyapal et al., 2026; Wright et al., 2023b). Collectively, these findings support the hypothesis that maternal sleep disruption may shape offspring outcomes through inflammatory pathways. However, the neurobiological mechanisms by which maternal sleep disruptions influence fetal neurodevelopment in utero remain poorly defined.

A shared consequence of inflammation and sleep deprivation is increased tryptophan degradation via the kynurenine pathway (KP) (Baratta et al., 2020; Notarangelo and Pocivavsek, 2017; Notarangelo and Schwarcz, 2021; Pocivavsek and Erhardt, 2024). The KP is initiated by the enzymes indoleamine 2,3-dioxygenase 1 (IDO1) and tryptophan 2,3-dioxygenase (TDO), which catalyze the conversion of tryptophan to kynurenine. Approximately 95% of dietary tryptophan undergoes catabolism through the KP, yielding nicotinamide adenine dinucleotide (NAD+), a ubiquitous cofactor and energy source, as well as neuroactive metabolites such as kynurenic acid (KYNA) and quinolinic acid (QUIN) (see **Figure 1**) (Pocivavsek et al., 2024). Notably, elevated KYNA levels are found in the cerebrospinal fluid and postmortem brain tissues of individuals with schizophrenia and bipolar disorder (Erhardt et al., 2001; Olsson et al., 2010; Sathyasaikumar et al., 2011; Schwarcz et al., 2001; Sellgren et al., 2019), populations characterized by high rates of sleep disturbances and chronic low-grade inflammation (Erhardt et al., 2017; Freeman and Waite, 2025; Halstead et al., 2023; Pocivavsek and Rowland, 2018). KYNA acts as an agonist at the G protein-coupled receptor (GPR) 35 (Wang et al., 2006), an orphan receptor, and the aryl hydrocarbon receptor (AhR) (DiNatale et al., 2010), which regulates immune signaling. KYNA also antagonizes the α7-nicotinic acetylcholine (α7nACh) (Hilmas et al., 2001) and N-methyl-D-aspartate (NMDA) (Stone, 1993) receptors, both critical for neurodevelopment and sleep regulation (Abreu-Villaça et al., 2011; Van Dongen, 2009; Wu et al., 2015). In preclinical studies, acute elevations in brain KYNA directly disrupt sleep and cognition (Chess et al., 2009; Pocivavsek et al., 2017; Pocivavsek et al., 2011; Rentschler et al., 2021), while prenatal KYNA elevations impair neurodevelopment and induce persistent biochemical and behavioral deficits in adult offspring (Buck et al., 2020; Hahn et al., 2018; Milosavljevic et al., 2023; Notarangelo and Pocivavsek, 2017; Pershing et al., 2015; Pershing et al., 2016; Pocivavsek et al., 2014; Pocivavsek et al., 2012; Rentschler et al., 2021; Wright et al., 2021). Together, these findings implicate KYNA as a potential mechanistic link between maternal sleep disruption and altered neurodevelopment.

**Figure 1:**
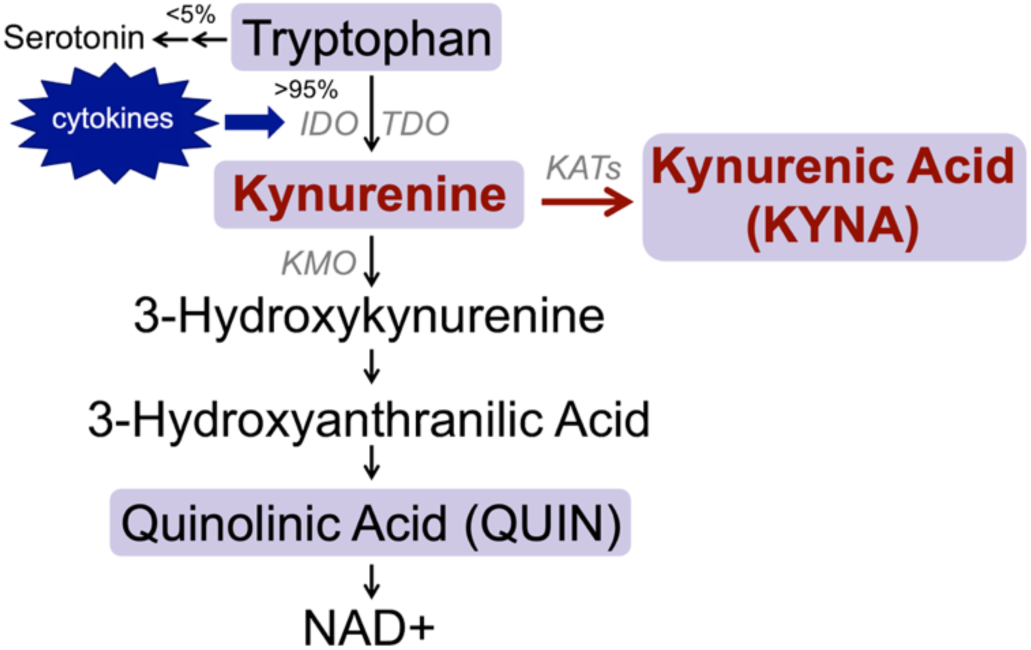
Schematic representation of tryptophan metabolism through the kynurenine pathway (KP). Tryptophan is degraded to kynurenine by *indoleamine dioxygenase* (*IDO*), which can be induced by inflammatory cytokines, or *tryptophan dioxygenase* (*TDO*). Kynurenine is subsequently degraded to kynurenic acid (KYNA) by *kynurenine aminotransferases* (*KAT*s) or to 3-hydroxykynurenine by *kynurenine 3-monooxygenase* (KMO), 3-hydroxyanthranilic acid, quinolinic acid (QUIN). The end product of the KP after QUIN is the energy-carrying molecule NAD+. Tryptophan-KP metabolites measured in the current study are highlighted in purple.

We previously demonstrated that acute maternal sleep deprivation (6 hours) increases placental inflammation and elevates fetal brain KYNA (Baratta et al., 2020). To expand and model clinically relevant chronic sleep disruption, we developed a sleep fragmentation (SleepFrag) paradigm and applied it during late gestation, from embryonic day (ED) 15 to ED 21, a period highly sensitive to inflammatory and KP perturbations (Cheslack-Postava and Brown, 2022; Pocivavsek et al., 2014). Maternal and fetal tissues were collected at ED 18 and ED 21. Despite the induction of peripheral inflammation with SleepFrag by ED 21, KP metabolites were not elevated in maternal plasma or brain, yet fetal brain KYNA was elevated. Then, in parallel, our embryonic kynurenine (EKyn) model was tested to increase prenatal KP metabolism via a maternal kynurenine-supplemented diet. EKyn treatment elevates fetal brain KYNA and produces sex-specific biochemical and behavioral deficits in adult offspring (Buck et al., 2020; Rentschler et al., 2021; Wright et al., 2021). Presently, we tested whether EKyn alters maternal sleep behavior and inflammation and investigated whether EKyn diet elevates KYNA to a differential degree in male and females prenatal tissues, including fetal brains. Whilst increasing maternal brain KYNA across regions, EKyn had minimal effects on maternal sleep behavior. However, EKyn produced a greater increase in KYNA in male compared to female fetal brains. Collectively, our models identify elevated fetal brain KYNA as a convergent molecular consequence of maternal inflammation and sleep disruption, with potential relevance for sex-specific neurobiological outcomes.

## 2. Methods

### 2.1. Animals

Adult (3-4 months) female and male Wistar rats were obtained from Charles River Laboratories. Female rats arrived naive or timed pregnant on embryonic day (ED) 3. For in-house timed pregnancy experiments, proven breeder male rats were used. All animals were singly housed upon arrival and allowed at least a week to acclimate to the animal facility before any experimentation. Animals received *ad libitum* food and water and were housed on a 12/12h light/dark cycle, where lights on corresponded to zeitgeber time (ZT) 0 and lights off to ZT 12. The animal facility was temperature-controlled and fully accredited by the American Association for the Accreditation of Laboratory Animal Care. All protocols were approved by the Institutional Animal Care and Use Committee at the University of South Carolina (USC) Floyd School of Medicine and were in accordance with the National Institutes of Health Guide for the Care and Use of Laboratory Animals.

### 2.2. Experimental Paradigms

#### 2.2.1. Experiment 1: Validation of Sleep Fragmentation (SleepFrag) protocol with custom sleep restriction chambers

Adult female rats (N = 9/group) were used to validate a novel sleep fragmentation (SleepFrag) protocol using custom automated sleep restriction chambers (Efird et al., 2024). Animals were implanted with EEG/EMG telemetry devices (described below), and sleep data were continuously acquired for 18 hours from ZT 12 (beginning of the dark phase) to ZT 6 (mid-light phase). Control subjects remained undisturbed in the home cage. For SleepFrag, sleep restriction chambers swept a bar across the cage floor to nudge the animals and disrupt sleep. The bar moved every 3 minutes from ZT 12 – 18, every 2 minutes from ZT 18 – 24, and every 1 minute from ZT 0 – 6. All animals had *ad libitum* access to food and water throughout the experiment.

#### 2.2.2. Experiment 2: SleepFrag during the last week of gestation

Maternal SleepFrag was conducted daily during the last week of gestation (ED 15 - ED 21). Control rats remained undisturbed in their home cages. Experimental rats underwent an automated sleep-restriction protocol for 18 hours daily (as described above). All animals were allowed undisturbed sleep from ZT 6-12 daily and had *ad libitum* access to food and water throughout the experiment. At ZT 6 on ED 18 (Control, N = 4, SleepFrag, N = 6) and ED 21 (Control, N = 4, SleepFrag, N = 6), dams were euthanized to harvest tissues for biochemical analysis (described below).

#### 2.2.3. Experiment 3: Embryonic Kynurenine (EKyn) Sleep Monitoring

Naive female rats (N=6/group) were implanted with EEG/EMG telemetry (described below). Following at least one week of surgical recovery, females in proestrus were housed overnight with a proven male breeder. Females were checked for a vaginal plug the next morning and returned to their home cage. In addition, female body weights were recorded on the day of pairing and 3 days post-pairing. A vaginal plug and/or 3-day weight gain greater than 10 g indicated a pregnancy. Females without a vaginal plug or significant weight gain post-pairing were re-paired with a different male the next time they entered proestrus. Care was taken to distribute male breeders among females and the experimental groups so that each male sired no more than one litter in each experimental group. Females were paired maximally twice before exclusion from the study. Embryonic kynurenine (EKyn; 100 mg kynurenine daily with 30 g wet mash) or control (ECon; 30 g wet mash) diet was fed daily during the last week of gestation (ED 15 to ED 21) (Pocivavsek et al., 2014). In pilot studies of dams implanted with EEG/EMG transmitters, we noted that labor started as early as ED 21. Thus, for the currently described sleep-monitoring experiments, sleep-wake behavior was evaluated on ED 20 for 24 h, beginning at ZT 0.

#### 2.2.4. Experiment 4: EKyn Diet Biochemical Analysis

Timed pregnant females (arrived on ED 3) were administered EKyn or ECon diet (described above) daily from ED 15. At ZT 6 on ED 21 (N = 4-8/group), dams were euthanized to harvest tissues for biochemical analysis (described below).

### 2.3. Chemicals

L-kynurenine sulfate salt (“kynurenine,” purity: 99.4%) was obtained from Sai Advantium (Hyderabad, India). For LC-MS/MS, and unless otherwise stated, chemicals were obtained from Sigma Aldrich (St. Louis, MO, USA). All other chemicals were obtained from various suppliers of the highest commercially available purity.

### 2.4. Sleep-Wake Behavior Monitoring

#### 2.4.1. Surgical procedure

Naive female Wistar rats were implanted intraperitoneally with radiotelemetry devices (PhysioTel HD-S02, Data Sciences International (DSI), St. Paul, MN, USA) to acquire electroencephalography (EEG) and electromyography (EMG) and monitor sleep-wake behavior as previously described (Pocivavsek et al., 2017). Under isoflurane anesthesia (3 – 5%), animals were placed in a stereotaxic frame (Stoelting Co., Wood Dale IL, USA). Carprofen (5 mg/kg, s.c.) was administered as an analgesic. EEG/EMG leads were threaded underneath the skin from a transverse incision into the abdominal cavity to reach a longitudinal incision along the midline of the head and neck. EEG leads were wrapped around two dental screws positioned over the frontal and parietal lobes (2.0 mm anterior/-1.5 mm lateral and 7.0 mm posterior/-1.5 mm lateral relative to bregma) and secured with acrylic dental cement. EMG leads were sutured into the cervical neck muscle approximately 1.0 mm apart. Incisions were closed with suture or wound clips. Animals were singly housed after surgery and allowed at least one week of surgical recovery.

#### 2.4.2. Sleep-wake data acquisition and analysis

Sleep-wake data were continuously acquired at a sampling rate of 500 Hz with Ponemah 6.10 software (DSI). Digitized signal data were scored offline with an automated Long Short-Term Memory (LSTM) machine learning algorithm (Smith et al., 2025) and NeuroScore 3.4 (DSI) in 10-s epochs as wake (low-amplitude, high-frequency EEG and high-amplitude EMG), non-rapid eye movement sleep (NREM; high-amplitude, low-frequency EEG and little to no EMG activity), or rapid eye movement sleep (REM; low-amplitude, high-frequency EEG with no EMG activity). All recordings initially classified by LSTM were reviewed in NeuroScore by human expert scorers.

### 2.5. Tissue Collection

On the day of tissue collection, animals were euthanized with CO_2_ asphyxiation at ZT 6. Whole maternal trunk blood was collected in tubes containing 25 μL K3-EDTA (0.15%) as an anticoagulant and centrifuged (1000 g, 10 minutes) to separate plasma. Litter sizes and the weight of each fetal body and placenta were recorded. Maternal brain, placentas, fetal livers, and fetal brains were rapidly removed, dissected on wet ice, and snap frozen on dry ice. Snips of fetal tails were collected for genetic sex determination (described below). Fetal tail snips were stored at −20°C. All other biological samples were stored at −80°C.

### 2.6. Determination of Fetal Sex

Biological sex was determined in ED 18 and ED 21 fetuses (also confirmed with visual sex determination, described below) by genotype assay for *Kdm5c* (X-chromosome-specific gene) and *Kdm5d* (Y-chromosome-specific gene), with slight modification from Dhakal and Soares 2017 (Dhakal and Soares, 2017). DNA was extracted from a tail snip by phenol chloroform extraction (Milosavljevic et al., 2025), and DNA concentration was measured via spectrophotometry (BioPhotometer Plus, Eppendorf, Enfield, CT, USA). DNA was amplified using polymerase chain reaction (PCR), forward primers for *Kdm5c* (TTTGTACGACTAGGCCCCAC) and *Kdm5d* (TTGGTGAGATGGCTGATTCC) sequences, and a common reverse primer (GTTTCTTAAACCGTCGCC: Thermo Fisher Scientific, Waltham, MA, USA). PCR reaction (20 μL per sample) contained 10 μL 2X master mix (Q5 hot start high-fidelity 2X Master Mix, New England Biolabs, Ipswich, MA, USA), 8 μL DNase/RNase-free distilled ultrapure water, 1 μL primer mix (1:1:1 primer mix from 10 μM stock), and 1 μL 25 ng DNA. Amplification was performed with an initial DNA denaturing step for 2 minutes at 95°C, followed by 35 cycles consisting of 30s denaturing at 95°C, 30s annealing at 55°C, 42s extension at 72°C, and a final extension for 7 minutes at 72°C (Eppendorf 5333 MasterCycler Thermal Cycler, Hamburg, Germany). PCR products were separated by agarose gel electrophoresis (MyGel Mini Electrophoresis System, Accuris Instruments, Edison, NJ, USA) at 110 volts for 50 minutes in 2% agarose gels containing 0.01% ethidium bromide based on size (250 bp, *Kdm5d* Y-chromosome gene and 692 bp, *Kdm5c* X-chromosome gene) and visualized using ultraviolet light (UVP High-Performance UV Transilluminators, Thermo Fisher Sceintific, Waltham, MA, USA).

In ED 21 samples, inspection of the distance between anus and external genitalia (anogenital distance) was used to determine biological sex (McCarthy, 2015). Male pups had a greater anogenital distance than female pups (**Supplemental Figure 1**). Sex determination by anogenital distance was confirmed by PCR as above.

### 2.7. Biochemical Analysis

#### 2.7.1. Corticosterone

Corticosterone was analyzed in plasma (6 μL diluted 1:40 v/v with steroid displacement reagent) with an enzyme-linked immunosorbent assay (ELISA), according to the manufacturer’s instructions (Enzo Life Sciences, Farmingdale, NY, USA). ELISA plates were read with a Synergy 2 Gen 5 microplate reader at 405 nm absorbance (BioTek Instruments Inc., Winooski, VT, USA).

#### 2.7.2. Cytokines

Cytokines were measured in maternal plasma (diluted 1:2 v/v in phospho-buffered saline, pH: 7.5) by a Rat Cytokine/Chemokine 27-Plex Discovery Assay Array (Eve Technologies, Calgary, AB, Canada). Selected cytokines were measured in placental samples using custom Bio-Plex Pro Rat Cytokine 1 5-plex Express Assays designed to detect IFN-γ, IL-1β, IL-6, IL-10, and TNF-α (Bio-Rad Laboratories Inc., Hercules, CA, USA) according to the manufacturer’s instructions. Briefly, tissues were thawed on wet ice and sonicated 1:6 w/v in lysis buffer (137 mM NaCl, 20 mM Tris- HCl, 10% Glycerol, 1% Tergitol-type NP40). Homogenates were centrifuged, the supernatant was collected, and protein content was determined using the Lowry method (LOWRY et al., 1951). Based on these values, aliquots containing 275 μg/125 μL of protein were prepared for Bio-Plex analysis and stored at −20°C until the day of each assay. Plates were read using a Bio-Rad Bio Plex 200 plate reader and Bio Plex Manager software. IL-10 and TNF-α were not detectable in our samples.

#### 2.7.3. Tryptophan, Kynurenine, and Kynurenic Acid (KYNA)

The contents of tryptophan, kynurenine or KYNA content in maternal plasma (diluted v/v in ultrapure water; 1:1000 for tryptophan, 1:10 for kynurenine and KYNA), maternal brain (sonicated 1:5 w/v in ultrapure water, final dilution 1:10 for kynurenine and KYNA), placenta (sonicated 1:10 w/v in ultrapure, final dilution 1:1000 for tryptophan, 1:50 for kynurenine, or 1:200 for KYNA), fetal liver (sonicated 1:5 w/v in ultrapure water, final dilution 1:1000 for tryptophan, 1:200 for KYNA), and fetal brain (sonicated 1:5 w/v in ultrapure water, final dilution 1:1000 for tryptophan, 1:10 for kynurenine and KYNA) were determined by ultra-high-performance liquid chromatography (UPLC). 80 μL of diluted sample was acidified with 20 μL of perchloric acid (6% saliva and plasma, 25% brain, liver, and placenta) and centrifuged (12,000 rpm, 10 minutes). 20 μL of the supernatant was isocratically eluted from a ReproSil-Pur C18 column (4mm x 100 mm; Dr. Maisch, GmbH, Ammerbuch, Germany), using a mobile phase containing 50 mM sodium acetate and 3-5% acetonitrile (pH adjusted to 6.2 with glacial acetic acid) at a flow rate of 0.5 mL/minute, and detected in the eluate with 500 mM zinc acetate delivered after the column with a flow rate of 0.1 mL/minute. In the eluate, tryptophan (excitation: 285 nm, emission: 365 nm), kynurenine (excitation: 365 nm emission: 480 nm), and KYNA (excitation 344 nm, emission 398 nm) were detected fluorometrically (Acquity UPLC H-Class PLUS System (Waters Corporation, Milford, MA, USA) with Empower 3 software (Waters Corporation) (Baratta et al., 2018b). Tissue protein content was determined in stock homogenate using the Lowry method (LOWRY et al., 1951).

#### 2.7.4. Quinolinic Acid (QUIN)

The content of QUIN in maternal plasma was determined by liquid chromatography tandem mass spectrometry (LC-MS/MS) (**Supplemental Figure 2**). 50 μL of plasma was added to 200 μL ice-cold methanol containing 10 pmol QUIN-d3 internal standard (MedChemExpress, Monmouth Junction, NJ, USA), calibrated to the weight of freshly prepared QUIN. The suspension was left on ice for 20 minutes and centrifuged (13,000 g, 10 minutes at 4°C). The supernatant was dried *in vacuo*. A QUIN standard curve ranging from 5 fmol (limit of detection) to 50 pmol was prepared. 10 pmol QUIN-d3 was added to each standard. The standard curve was linear across all standards. QUIN plasma levels ranged from 5 fmol to 2.5 pmol. Solid Phase Extraction (SPE) was performed on both standards and plasma samples using 1 cc 18 SPE cartridges (Waters Corporation) with 0.1% trifluoroacetic acid/40% methanol as the eluent.

The eluents were dried *in vacuo* prior to derivatization. 25 μL of ethanol/acetyl chloride (100:20 v/v, mixed on ice) was added to each sample. The samples were vortexed thoroughly, incubated (55°C, 2 h), centrifuged (10 s), and dried completely *in vacuo* (Cox et al., 2024). The reaction products were resuspended in 200 μL acetonitrile/water (20:80 v/v) and transferred to autosampler vials for LC-MS/MS analysis. LC separation was performed on a Thermo Vanquish Flex liquid chromatography system (ThermoFisher Scientific) using a Waters XBridge C18 reverse-phase column (2.1 mm x 100 mm, 3.5 μm particles; Waters Corporation) maintained at 40°C. Following a 3 μL sample injection, a binary gradient elution was performed. Solvent A consisted of water with 0.1% formic acid and solvent B contained acetonitrile with 0.1% formic acid. The separation began with the column equilibrated at 10% B for 1.5 minutes then ramped to 95% B at 10 minutes and held there until 15 minutes. The system then returned to 10% B for 8 minutes. The column flow rate was 0.2 μL/minute. Single injections of QUIN standards confirmed their separation and retention times.

Positive ion electrospray mass spectra were acquired on a Thermo Q-Exactive HF- X Quadrupole-Orbitrap performing parallel reaction monitoring (PRM). Precursor ions (224 and 227 Da) were isolated by the quadrupoles using a 0.5 m/z isolation window and fragmented in the higher-energy collisional dissociation (HCD) cell using stepped collision energy (CE) (15,25,35 eV); the automatic gain control (AGC) target was 1E6 and maximum ion trap (IT) was set at 150 ms. The Orbitrap resolution used for PRM was 30,000. The source capillary temperature was 275°C, other MS source settings were sheath gas flow: 45; auxiliary gas flow: 10; sweep gas flow: 2; spray voltage: 3.5 kV; funnel RF level: 40; auxiliary gas temperature: 400°C. XCalibur™ 4.2 software (ThermoFisher) was used to construct extracted ion chromatograms of the transitions 224→150 (QUIN), and 227→153 (QUIN-d3). The area of the QUIN peak was normalized to the area of the QUIN-d3 internal standard to obtain peak area ratios. The mass of QUIN in the plasma samples was normalized to the plasma volume to calculate QUIN concentration.

### 2.8. Data Handling and Statistics

Each maternal and fetal sample was an independent biological data point. To minimize the contribution of individual litters, placental and fetal samples were handled such that randomized samples (N=1-4/sex/litter) were analyzed in all biochemical experiments. Maternal plasma inflammatory markers were log2-transformed for heat map visualizations to highlight fold-changes between experimental groups and manage the wide range of concentrations. All statistical analyses were conducted on raw values (pg/mL). EGF (N=1), GRO (N=1), IL-1β (N=1), IFN-γ (N=1), MCP-1 (N=3), TNF-α (N=4), IL-10 (N=2), were not detectable in ED21 SleepFrag maternal plasma. Statistical analysis was not conducted for inflammatory markers with < 3 detectable samples. TNF-α and IL-10 were below the limit of detection in placental samples and were excluded from analysis. Across all measures, outliers were identified using ROUT analysis, with Q = 1%, and excluded from analysis, data are expressed as mean ± SEM, and statistical analyses were performed using GraphPad Prism 11.0 software (GraphPad Software, La Jolla, CA, USA) with statistical significance defined as *p* < 0.05. Significant two-way ANOVA results were followed up by Fisher’s LSD post hoc tests. A supplemental statistical file is included to present all ANOVA results.

#### 2.8.1. Experiment 1: Validation of SleepFrag protocol

SleepFrag sleep architecture data were analyzed across the 18-hr period using Mann-Whitney tests for each vigilant state (Wake, NREM sleep, REM sleep). Control data represent the average of four days of *ad libitum* sleep-wake behavior recording.

#### 2.8.2. Experiment 2: Gestational SleepFrag Biochemical Analyses

Maternal plasma cytokine levels were first analyzed using Mann-Whitney tests, with each ED and cytokine analyzed separately. Cytokines with significant Mann-Whitney tests, along with all other maternal biochemistry, were assessed using two-way ANOVAs with ED and treatment as between-subjects factors. Maternal brain analytes were analyzed separately by brain region. Fetal biochemistry data were analyzed using three-way ANOVAs with ED, SleepFrag, and sex as between-subject factors. Given the effect of SleepFrag at ED 21 but not at ED 18 in our analyses, and our hypothesis that SleepFrag and biological sex will significantly influence our measured outcomes, we also conducted two-way ANOVAs stratified by ED.

#### 2.8.3. Experiment 3: EKyn Diet Sleep Monitoring

ECon/EKyn sleep duration was analyzed in 12-hr bins (ZT 0-12, light phase, or ZT 12-24, dark phase) by three-way repeated measures (RM) ANOVA with treatment as a between-subject factor and light phase and vigilant state as within-subject factors and two-way RM ANOVA with treatment as a between-subject factor and light phase as a within-subject factor. Total sleep duration was calculated as the sum of REM sleep and NREM sleep.

#### 2.8.4. Experiment 4: EKyn Diet Biochemical Analyses

ECon/EKyn maternal biochemistry data were analyzed by Mann-Whitney tests. Maternal brain analytes were analyzed separately by brain region. Fetal data were analyzed by two-way ANOVA with treatment and sex as between-subject factors.

## 3. Results

### 3.1. Sleep-wake architecture is altered with sleep fragmentation (SleepFrag) protocol

We first characterized the effects of SleepFrag (**Figure 2A**) on sleep-wake architecture in female rats across 18 hours. SleepFrag significantly increased time spent awake (**Figure 2D**: +43%, *p* < 0.0001), and reduced NREM sleep (−58%, *p* < 0.0001) and REM sleep (−94%, *p* < 0.0001). SleepFrag increased the number of vigilant state bouts (**Figure 2G**; wake: +56%, *p* < 0.001; NREM sleep: + 34%, *p* = 0.077) except for REM sleep, which was nearly abolished (−89%, *p* < 0.0001). SleepFrag led to sustained bouts of wakefulness (**Figure 2J**: +112%, *p* = 0.0503) and reduced the average bout duration of NREM sleep (−68%, *p* < 0.0001) and REM sleep (−45%, *p* < 0.001).

**Figure 2:**
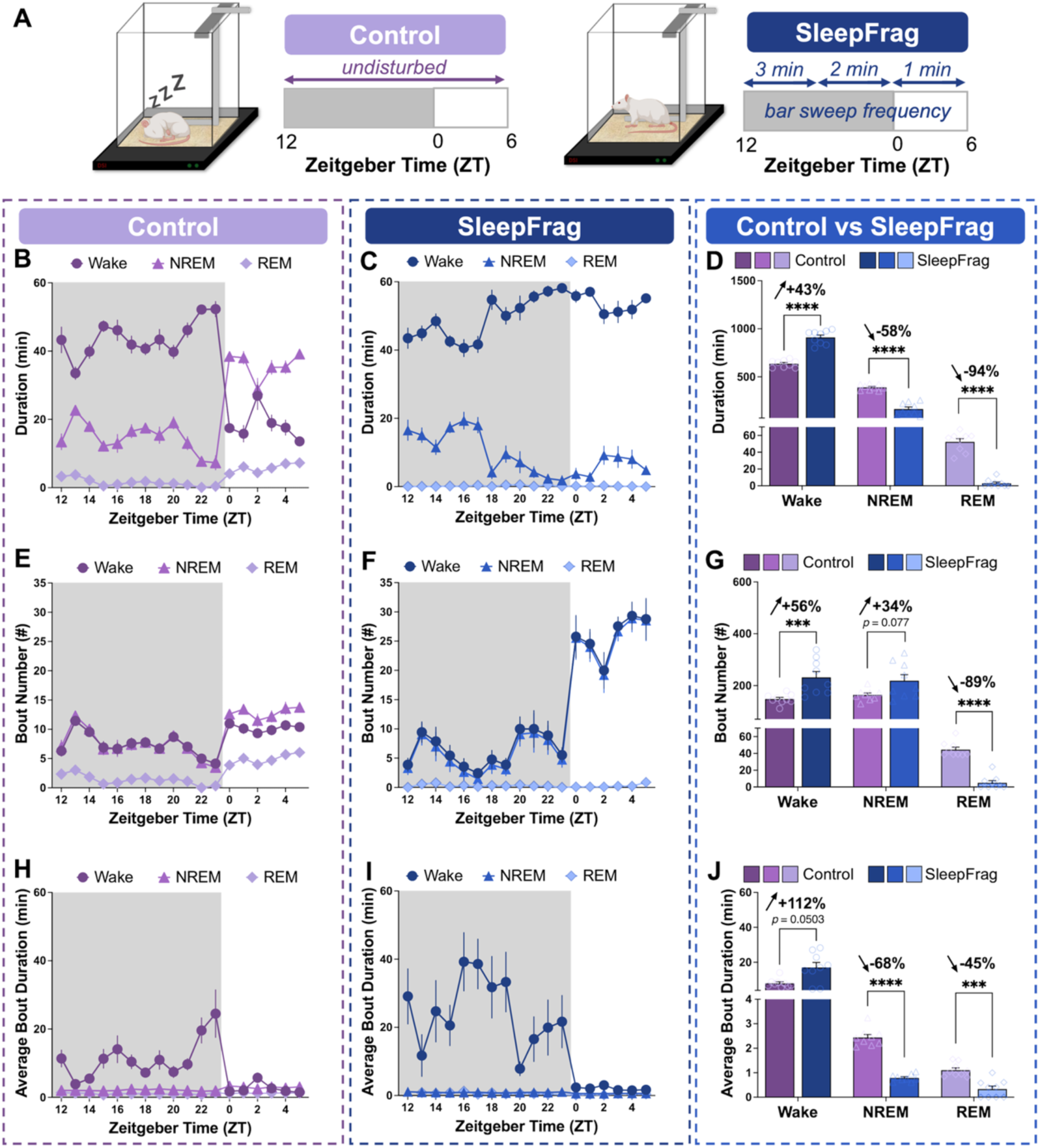
Sleep fragmentation (SleepFrag) protocol dramatically reduces sleep duration and increases the frequency of vigilant-state bouts. **(A)** Visual representation of experimental conditions. Control rats remained undisturbed for 18 hours from Zeitgeber time (ZT) 12 to ZT 6, spanning the entire dark phase and half of the subsequent light phase. Data from control rats are the average of four days of undisturbed sleep-wake behavior recording. SleepFrag rats were subjected to 18 hours of SleepFrag from ZT 12 to ZT 6 by a bar sweeping across the bottom of the home cage every 3 minutes (ZT 12-18), 2 minutes (ZT 18-24), or 1 minute (ZT 0-6). **(B/C)** Vigilant state durations in 1-hour bins in Control and SleepFrag. **(D)** Sum of vigilant state durations for experiment duration (18 hours). **(E/F)** Vigilant state bout numbers in 1-hour bins in Control and SleepFrag. **(G)** Sum of vigilant state bout numbers for experiment duration (18 hours). **(H/I)** Vigilant state average bout duration in 1-hour bins in Control and SleepFrag. **(J)** Average of vigilant-state bout duration for the experiment duration (18 hours). Data are mean ± SEM, with individual data points shown in bar graphs. Mann-Whitney tests: *** *p* < 0.001, **** *p* < 0.0001. N = 9/group.

### 3.2. Maternal SleepFrag increases peripheral inflammation during pregnancy

SleepFrag was induced during pregnancy from ED 15 to ED 21, with tissue collection occurring after 3 days on ED 18 or after 6 days on ED 21 (**Figure 3A**). Maternal plasma corticosterone and inflammatory markers were assessed to measure the impact of SleepFrag on the stress response and immune system activation. At both ED 18 and ED 21, maternal plasma corticosterone was not elevated compared to control (**Figure 3B**). Yet several cytokines were significantly elevated in ED 21 SleepFrag maternal plasma compared to control (**Figure 3C**, individual *p*-values are listed in **Supplemental Table 1**). We identified IFN-γ, IL-1β, IL-6, and IL-17A as cytokines previously implicated in KP activation and associated with offspring neurodevelopmental disorders (**Figure 3D-G**)(Badawy, 2017; Baratta et al., 2020; Choi et al., 2016; Han et al., 2021; Otero and Antonson, 2022; Wu et al., 2017). All remaining significant results are displayed individually in **Supplemental Figure 3**. Analysis of IFN-γ revealed a significant interaction between SleepFrag and ED (F_1, 14_ = 15.82, *p* < 0.01) and a main effect of SleepFrag (F_1, 14_ = 6.521, *p* < 0.05)(**Figure 3D**). SleepFrag significantly elevated IFN-γ at ED 21 (*p* < 0.001). IL-1β levels were impacted by ED (F_1, 15_ = 7.114, *p* < 0.05)(**Figure 3E**). IL-6, elevated in SleepFrag dams compared to controls at ED 21 (*p* < 0.01), showed a significant ED x SleepFrag interaction (F_1, 16_ = 10.05, *p* < 0.01) and a main effect of SleepFrag (F_1, 16_ = 5.246, *p* < 0.05)(**Figure 3F**). IL-17A levels were changed across ED, noted with an ED x SleepFrag interaction (F_1, 16_ = 10.64, *p* < 0.01)(**Figure 3G**). Compared to ED 18, IL-17A levels at ED 21 were lower in control plasma (*p* < 0.05) and higher in SleepFrag plasma (*p* < 0.05). At ED21, SleepFrag elevated IL-17A levels compared to control (*p* < 0.01)

**Figure 3:**
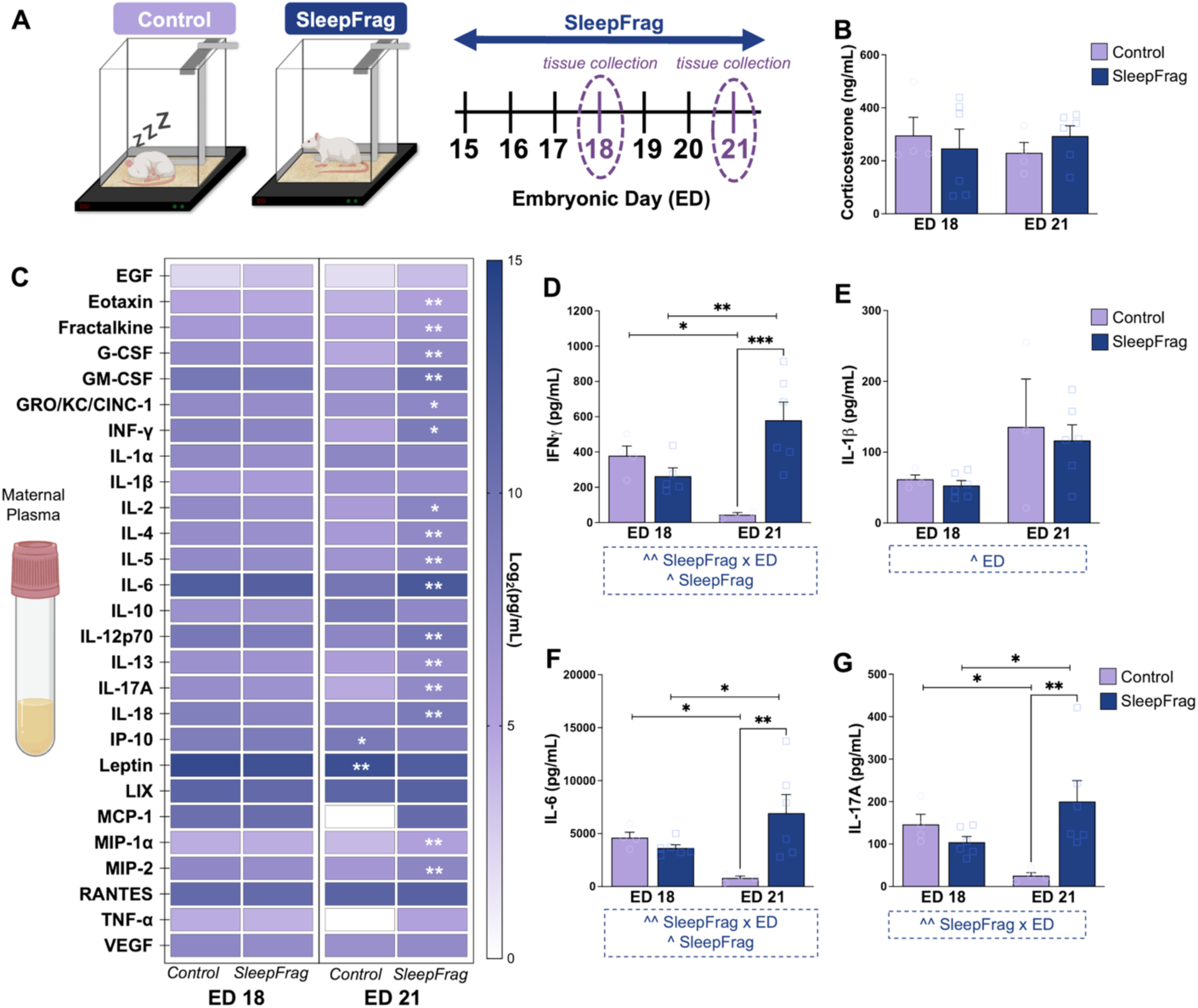
In dams, SleepFrag increases inflammatory markers in maternal plasma during pregnancy. **(A)** Schematic of experimental paradigm. From embryonic day (ED) 15 to ED 21, SleepFrag was induced daily for 18 hours from Zeitgeber time (ZT) 12 to ZT 6, spanning the entire dark phase and half of the subsequent light phase. Control subjects remained undisturbed in the home cage. Tissues were collected at ZT 6 on ED 18 and ED 21. Maternal plasma **(B)** corticosterone, **(C)** 27-plex cytokine discovery panel, **(D)** IFN-γ, **(E)** IL-1β, **(F)** IL-6, **(G)** IL-17A. Data are mean ± SEM with individual data points shown in bar graphs. Two-way ANOVA conducted for Panels B, D, E, F, G with significant ANOVA results displayed in blue boxed annotations below figure panels: ^ *p* < 0.05, ^^ *p* < 0.01, and post-hoc Fisher’s LSD: * *p* < 0.05, ** *p* < 0.01, *** *p* < 0.001. Mann-Whitney test conducted for Panel C showing control versus SleepFrag at ED 18 or ED 21: * *p* < 0.05, ** *p* < 0.01. N = 3-6 dams/group.

### 3.3. Maternal SleepFrag does not elevate kynurenine pathway (KP) metabolites in the maternal compartment

Given that several pro-inflammatory cytokines were elevated following SleepFrag, we tested whether this would elevate KP metabolites. We first assessed tryptophan and its metabolites in maternal plasma. ED (F_1, 17_ = 26.57, *p* < 0.0001) but not SleepFrag impacted maternal plasma tryptophan (**Figure 4A**) while maternal plasma kynurenine remained unaffected by ED or SleepFrag (**Figure 4B**). We then evaluated quinolinic acid (QUIN), the alternate branch of KP metabolism, and determined a significant interaction between ED and SleepFrag in maternal plasma QUIN levels (F_1, 16_ = 16.08, *p* < 0.01), as well as a main effects of ED (F_1, 16_ = 19.27, *p* < 0.001) and SleepFrag (F_1, 16_ = 29.19, *p* < 0.0001) (**Figure 4C**). Plasma QUIN in maternal controls was higher at ED 21 than at ED 18 (*p* < 0.0001), while QUIN levels remained unchanged across ED with SleepFrag. Thus, QUIN levels were significantly lower at ED 21 in SleepFrag compared to controls (*p* < 0.0001). Alternatively, we found that maternal plasma KYNA levels were significantly affected by an ED x SleepFrag interaction (F_1, 16_ = 5.223, *p* < 0.05) (**Figure 4D**), wherein at ED 21, maternal plasma KYNA levels were reduced after SleepFrag (*p* < 0.05).

**Figure 4:**
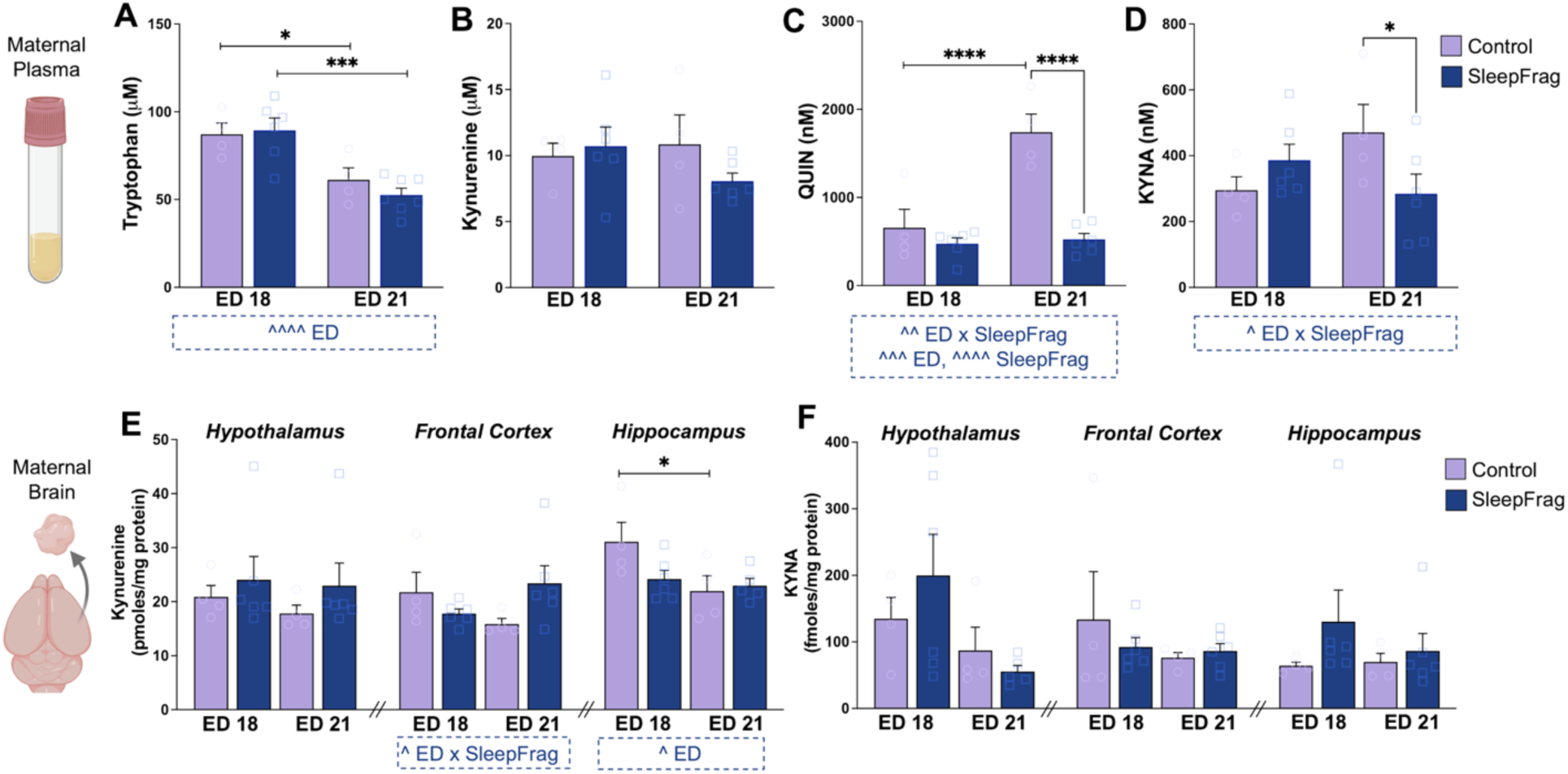
In dams, SleepFrag reduces peripheral kynurenine pathway metabolism but does not alter maternal brain KYNA. From embryonic day (ED) 15 to ED 21, SleepFrag was induced daily for 18 hours from Zeitgeber time (ZT) 12 to ZT 6, spanning the entire dark phase and half of the subsequent light phase. Control subjects remained undisturbed in the home cage. Tissues were collected at ZT 6 on ED 18 and ED 21. Maternal plasma **(A)** tryptophan, **(B)** kynurenine, **(C)** QUIN, **(D)** KYNA. Maternal brain **(E)** kynurenine, **(F)** KYNA. All data are mean ± SEM with individual data points shown. Significant two-way ANOVA results are displayed in blue boxed annotations below figure panels: ^ *p* < 0.05, ^^ *p* < 0.01, ^^^ *p* < 0.001, ^^^^ *p* < 0.0001. Fisher’s LSD post-hoc: * *p* < 0.05, *** *p* < 0.001, **** *p* < 0.0001. N = 4-6 dams/group.

To approximate the enzymatic conversion of (i) tryptophan to kynurenine via IDO and TDO, (ii) kynurenine to KYNA via KATs, (iii) kynurenine to QUIN via the KMO branch of the pathway, and (iv) the balance between the respective KYNA and KMO branches, we calculated the (i) kynurenine to tryptophan (kynurenine/tryptophan), (ii) KYNA to kynurenine (KYNA/kynurenine), (iii) QUIN to kynurenine (QUIN/kynurenine), and (iv) KYNA to QUIN (KYNA/QUIN) ratios. Of note, the QUIN/kynurenine ratio was reduced while the KYNA/QUIN ratio was increased with SleepFrag (**Supplemental Figure 4**), suggesting greater activity to KYNA production in the KP over QUIN production. Thus, subsequent tissue analyses were restricted to assessing tryptophan, kynurenine, and KYNA content.

To investigate the impact of SleepFrag on the maternal brain, we assessed kynurenine and KYNA in the hypothalamus, frontal cortex, and hippocampus, brain regions that undergo dynamic changes during pregnancy and are critical for homeostatic regulation, maternal behavior, and cognition (Chechko et al., 2022). Across these brain areas, SleepFrag did not significantly change maternal brain kynurenine (**Figure 4E**) or KYNA (**Figure 4F**) levels. Of note, there was a strong trend toward a main effect of ED on KYNA in the hypothalamus (F_1, 7_ = 5.108, *p* = 0.0583), mainly driven by elevated KYNA levels at ED 18 and reduced KYNA levels at ED 21 in SleepFrag maternal brains.

### 3.4. Maternal SleepFrag elevates kynurenine pathway (KP) metabolism in the fetal compartment and elevates KYNA in the fetal brain

Next, we sought to characterize the impact of SleepFrag in the fetal compartment. As published studies suggest sex-specific responses to prenatal insults, neurodevelopmental disorders, and KP alterations (Bale, 2016; Buck et al., 2020; Hall et al., 2023; Milosavljevic et al., 2025; Rentschler et al., 2021; Wright et al., 2021), we differentiated our prenatal analyses by biological sex. First, we confirmed that SleepFrag did not affect broad developmental characteristics of the placentas or fetuses (**Supplemental Figure 5**) and found mildly reduced body weight in ED 21 SleepFrag fetuses compared to controls, but no differences in placental weight, placental efficiency, litter size, or the sex ratio of viable fetuses between experimental groups.

The placenta, an active endocrine and immune organ during gestation, serves as a key mediator of maternal immune responsivity and fetal brain neurodevelopment (Kramer et al., 2023). In the placenta, we measured levels of IFN-γ, IL-1β, and IL-6, immune factors associated with KP activation (Badawy, 2017). The impact of SleepFrag, ED, and biological sex was evaluated for each placental cytokine. IFN-γ (**Figure 5A**) and IL-1β (**Figure 5B**) were not significantly influenced by any factor. IL-6 was influenced by a significant SleepFrag x ED interaction (F_1, 33_ = 8.820, *p* < 0.01) (**Figure 5C**). At ED 21, SleepFrag male placentas had elevated IL-6 levels compared to control (*p* < 0.01). Female placental IL-6 levels were not different from control at either time point.

**Figure 5:**
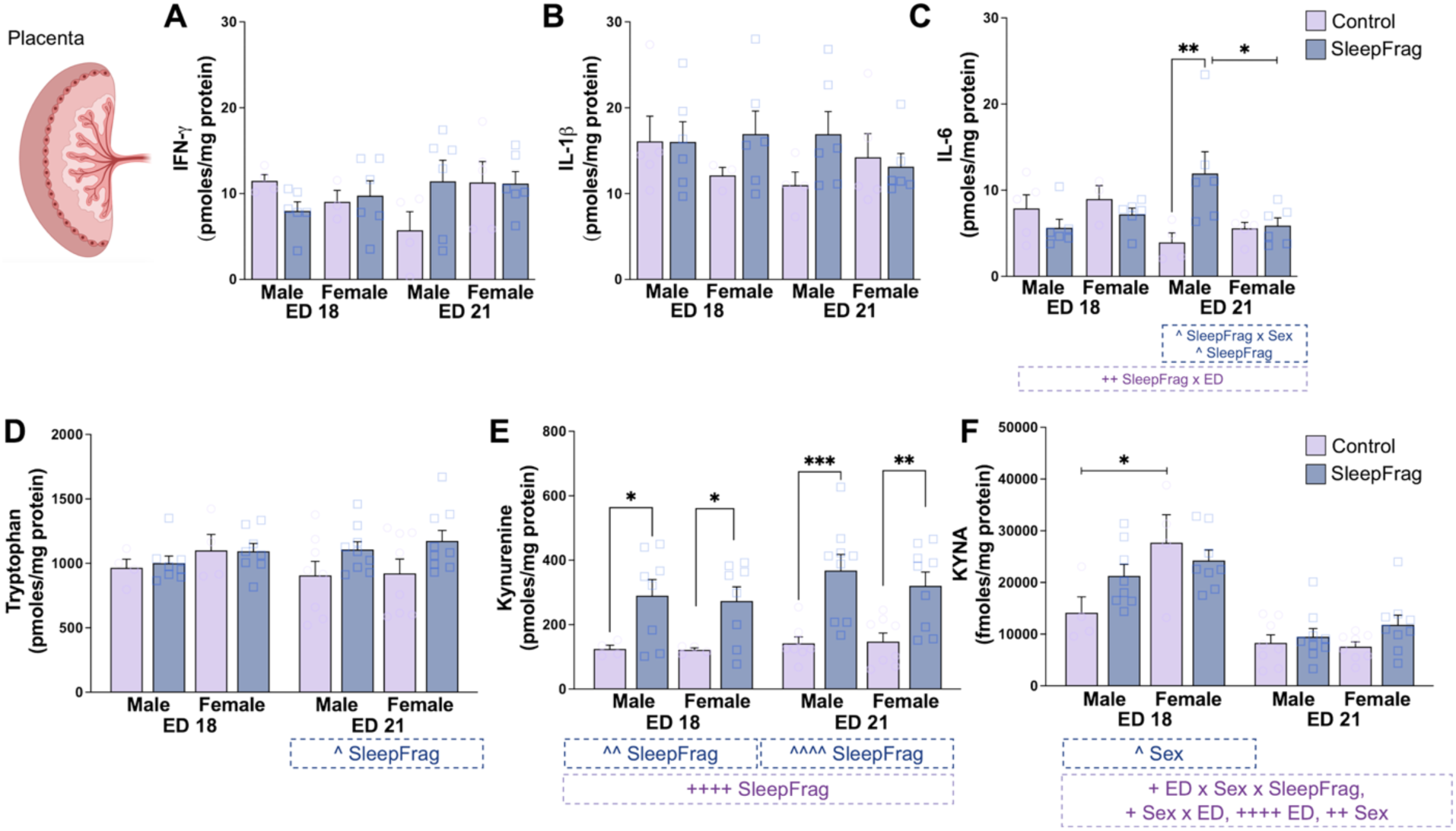
In placental tissue, SleepFrag increases IL-6 and tryptophan metabolism to heighten kynurenine and KYNA formation. From embryonic day (ED) 15 to ED 21, SleepFrag was induced daily for 18 hours from Zeitgeber time (ZT) 12 to ZT 6, spanning the entire dark phase and half of the subsequent light phase. Control subjects remained undisturbed in the home cage. Tissues were collected at ZT 6 on ED 18 and ED 21. Placental **(A)** IFN-γ, **(B)** IL-1β, and **(C)** IL-6, **(D)** tryptophan, **(E)** kynurenine, and **(F)** KYNA. Data are mean ± SEM with individual data points shown. Significant three-way ANOVA results are displayed in purple boxed annotations below each figure panel: + *p* < 0.05, ++ *p* < 0.01, ++++ *p* < 0.0001. Significant two-way ANOVA results, separated by ED, are displayed in blue boxed annotations below each figure panel: ^ *p* < 0.05, ^^ *p* < 0.01, ^^^^ *p* < 0.0001. Fisher’s LSD post-hoc: * *p* < 0.05, ** *p* < 0.01, *** *p* < 0.001. N = 3-9 pups/group, N = 3-6 dams/group.

Placental metabolite analysis revealed that SleepFrag induced higher tryptophan levels at ED 21 (F_1, 30_ = 6.229, *p* < 0.05) compared to controls, regardless of sex (male: +1.3-fold, *p* < 0.001; female: + 1.2-fold, *p* < 0.01) (**Figure 5D**). SleepFrag also influenced kynurenine levels (**Figure 5E**), noted as elevated levels of kynurenine at ED 18 (main effect SleepFrag: F_1, 20_ = 10.83, *p* < 0.01) and ED 21 (main effect SleepFrag: F_1, 30_ = 27.87, *p* < 0.0001) compared to controls in both male and female placentas (ED 18 male: +2.3-fold, *p* < 0.05, female: +2.3-fold, *p* < 0.05; ED 21 male: +3-fold, *p* < 0.001, female: +2.2-fold, *p* < 0.01). Interactions between ED x Sex x SleepFrag (F_1, 50_ = 4.861, *p* < 0.05) and Sex x ED (F_1, 50_ = 5.840, *p* < 0.05) impacted placental KYNA levels, as well as main effects of ED (F_1, 50_ = 65.68, *p* < 0.0001) and Sex (F_1, 50_ = 8.623, *p* < 0.01) (**Figure 5F**). When assessed at either ED, placental KYNA levels were not elevated with SleepFrag.

To understand if placental KP alterations are related to changes in the fetus, we next investigated metabolites in the fetal liver and brain. Because the liver has high enzymatic activity that rapidly converts kynurenine into its downstream metabolites (Ceresoli-Borroni and Schwarcz, 2000; Notarangelo et al., 2019), we analyzed only tryptophan and KYNA. Across treatment conditions, higher liver tryptophan (main effect ED: F_1, 54_ = 66.98, *p* < 0.0001) and KYNA (main effect ED: F_1, 53_ = 169.0, *p* < 0.0001) were found at ED 21 compared to at ED 18 (**Supplemental Figure 6**). Fetal liver KYNA was impacted by a significant SleepFrag x ED x Sex interaction (F_1,53_ = 4.562, *p* = 0.0373). At ED 21, we found a SleepFrag x Sex interaction (F_1, 29_ = 4.621, *p* = 0.0401) wherein compared to control, SleepFrag lowered male fetal liver KYNA and elevated female liver KYNA.

Tryptophan, kynurenine, and KYNA were measured in the fetal brain. A significant ED x SleepFrag interaction (F_1, 61_ = 8.911, *p* < 0.01) and main effects of ED (F_1, 61_ = 36.53, *p* < 0.0001) and SleepFrag (F_1, 61_ = 12.54, *p* < 0.001) impacted fetal brain tryptophan (**Figure 6A**). At ED 21, SleepFrag (main effect F_1, 30_ = 16.89, *p* < 0.001) elevated fetal brain tryptophan compared to control in both sexes (male: +1.5-fold, *p* < 0.01, female: +1.5-fold, *p* < 0.05). Kynurenine was likewise influenced by a main effect of SleepFrag (F_1, 60_ = 9.812, *p* < 0.01) (**Figure 6B**). At ED 21, kynurenine levels were higher following SleepFrag (main effect: F_1, 29_ = 15.53, *p* < 0.001) in both male (+1.3-fold, *p* < 0.05) and female (+1.5-fold *p* < 0.01) fetal brains. Of particular interest, fetal brain KYNA was significantly influenced by an interaction between SleepFrag and ED (F_1, 58_ = 4.242, *p* = 0.0439), a main effect of ED (F_1, 58_ = 31.82, *p* < 0.0001) (**Figure 6C**). At ED 21, KYNA was elevated after SleepFrag (main effect: F_1, 28_ = 5.091, *p* < 0.05). Post-hoc analysis revealed that this elevation was driven by a more prominent increase in male fetal brains (male: +1.6-fold, *p* = 0.068, female: +1.3-fold, *p* = 0.21).

**Figure 6:**
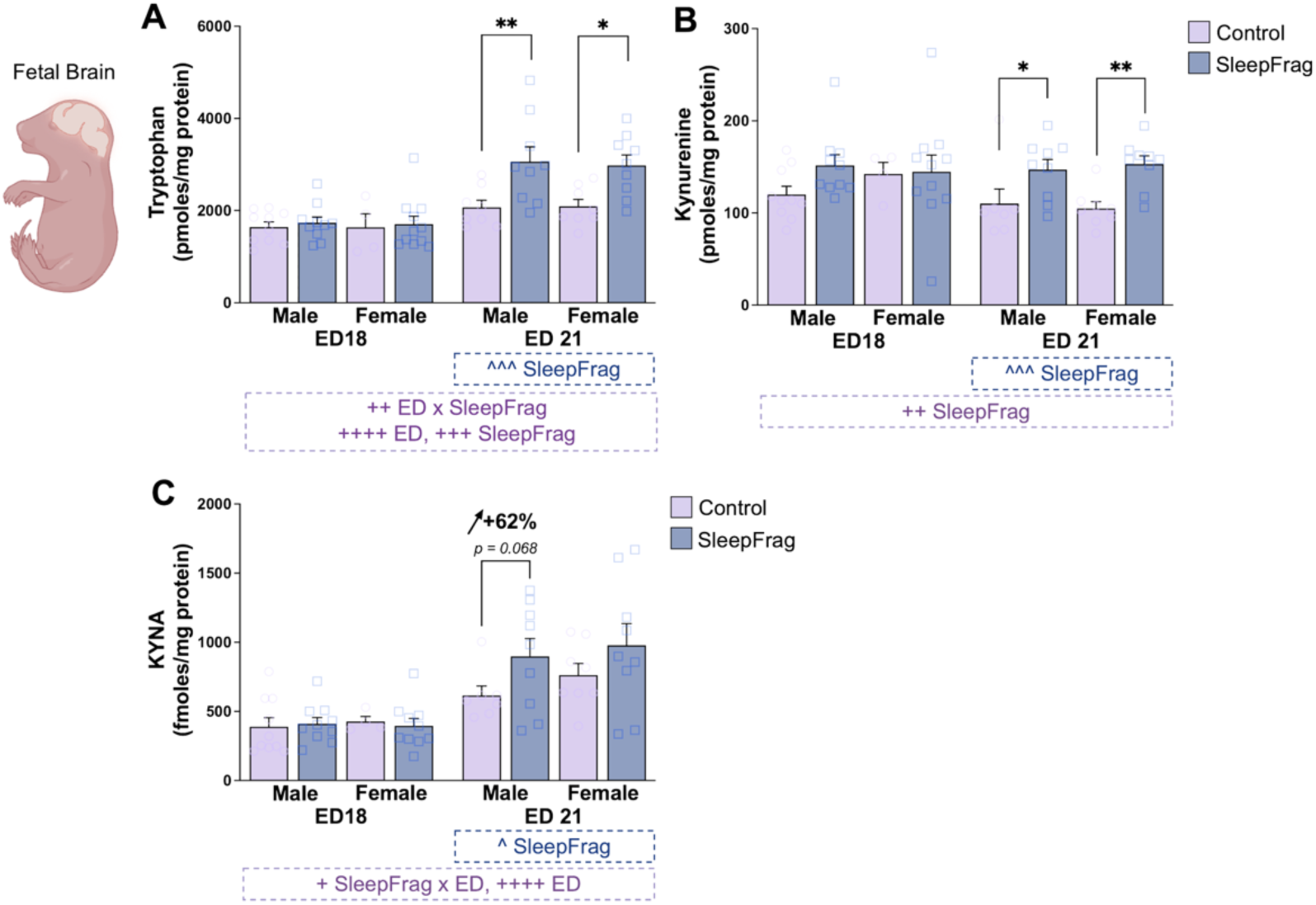
In the fetal brain, SleepFrag alters tryptophan metabolism to heighten kynurenine and KYNA formation. From embryonic day (ED) 15 to ED 21, SleepFrag was induced daily for 18 hours from Zeitgeber time (ZT) 12 to ZT 6, spanning the entire dark phase and half of the subsequent light phase. Control subjects remained undisturbed in the home cage. Tissues were collected at ZT 6 on ED 18 and ED 21. Fetal brain **(A)** tryptophan, **(B)** kynurenine, and **(C)** KYNA. Data are mean ± SEM with individual data points shown. Significant three-way ANOVA results are displayed in purple annotations below each figure panel: + *p* < 0.05, ++ *p* < 0.01, ++++ *p* < 0.0001. Significant two-way ANOVA results, separated by ED, are displayed in blue boxed annotations below each figure panel: ^ *p* < 0.05, ^^^ *p* < 0.001. Fisher’s LSD post-hoc: * *p* < 0.05, ** *p* < 0.01. N = 4-10 pups/group, N = 4-6 dams/group.

To approximate the enzymatic conversion of tryptophan to kynurenine via IDO1 and TDO or the kynurenine to KYNA conversion via KATs, we calculated the kynurenine/tryptophan and KYNA/kynurenine ratios in placenta and fetal brain (**Supplemental Figure 7).** Notably, SleepFrag elevated the kynurenine/tryptophan ratio in the placentas of both sexes at ED 18 and ED 21, while at ED 21, only the male fetal brain KYNA/kynurenine ratio increased compared to control.

### 3.5. Embryonic kynurenine (EKyn) diet does not cause major maternal sleep disruption

During the final week of gestation, maternal SleepFrag markedly disrupted sleep and revealed fetal brain vulnerability to KYNA elevation. To more precisely simulate the effects of gestational KYNA increases, the EKyn model increases KP metabolism during neurodevelopment by feeding dams a kynurenine-supplemented diet (Pocivavsek et al., 2014). We currently expanded EKyn diet studies to focus on maternal sleep behavior at the end of pregnancy, maternal plasma inflammatory markers, and biochemical consequences in maternal and sex-specific fetal compartments. Like the SleepFrag protocol, the EKyn model and counterpart control (ECon) were implemented during the last week of gestation, ED 15 to ED 22.

We have previously linked elevated brain KYNA to disrupted sleep architecture and reduced REM sleep (Milosavljevic et al., 2025; Pocivavsek et al., 2017; Rentschler et al., 2024). Notably, in nulliparous females, acute elevation of KYNA with systemic kynurenine treatment resulted in transient sleep changes (Rentschler et al., 2024), whereas the impact of EKyn diet on maternal sleep during pregnancy has been unexplored. Here, we determined only a subtle difference in sleep-wake behavior on ED 20 between ECon and EKyn dams (**Figure 7A**). Wake and total sleep duration were significantly impacted by a three-way interaction between EKyn x vigilant state x light phase (F_1, 18_ = 1025, *p* < 0.01) (**Figure 7B**). In the light phase, maternal sleep and awake duration were unaffected by EKyn diet. Yet during the dark phase, EKyn dams spent more time awake (*p* < 0.05) and less time asleep (*p* < 0.05) than ECon (EKyn x vigilant state interaction: F_1, 9_ = 5.5, *p* < 0.05). We also assessed relative cage activity and found main effects of EKyn (F_1, 9_ = 6.128, *p* < 0.05) and light phase (F_1, 8_ = 66.07, *p* < 0.0001). Importantly, EKyn dams were more active during the dark phase than ECon (*p* < 0.05), suggesting less quiet wake behavior typically observed in pregnant rodents (**Figure 7C**)(Komiya et al., 2018; Sivadas et al., 2017). Analysis of sleep-wake architecture by vigilant state yielded no significant post-hoc differences between EKyn and ECon dams (**Supplemental Figure 8**).

**Figure 7:**
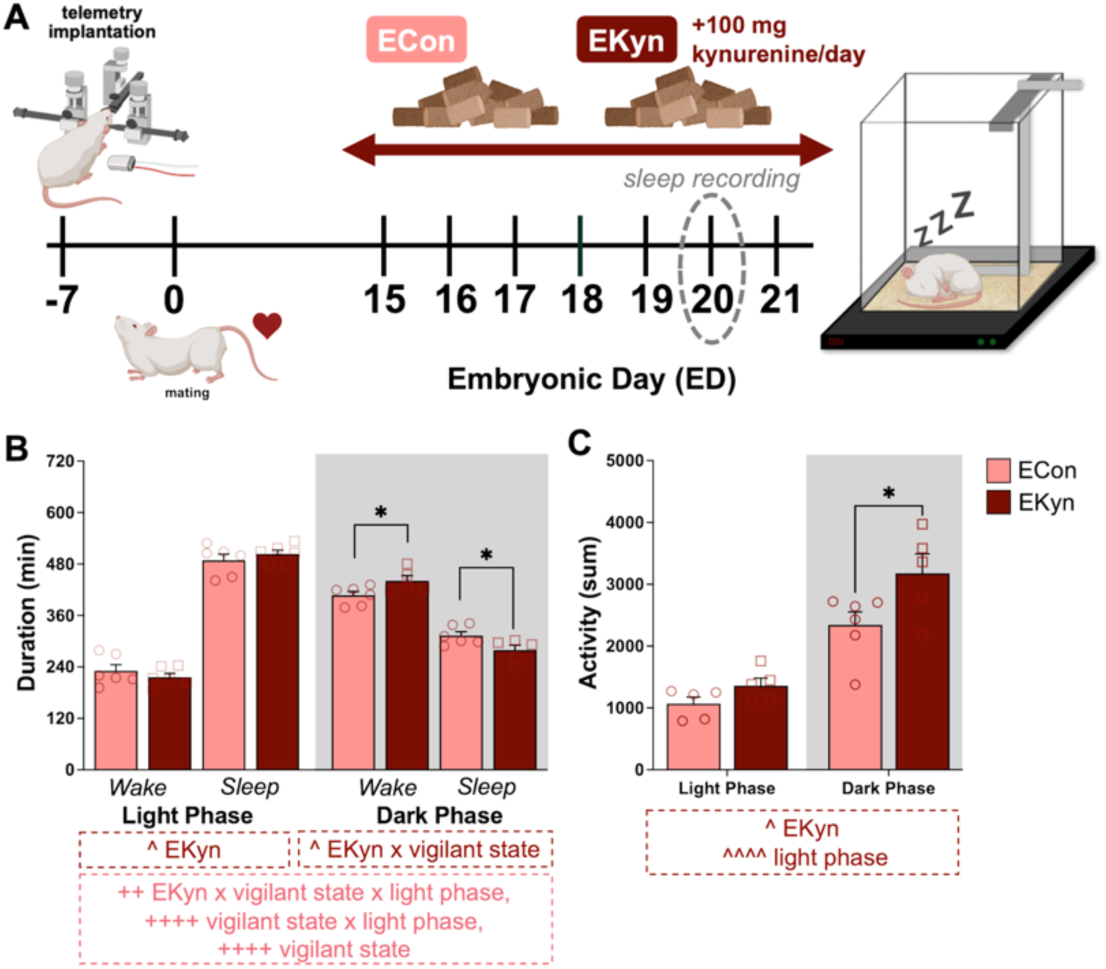
In dams, embryonic kynurenine (EKyn) diet increases dark phase wake duration and activity, while total sleep time decreases. **(A)** Schematic of experimental timeline. From embryonic day (ED) 15 to ED 21, pregnant females, who were implanted with radiotelemetry devices to obtain EEG/EMG data, received embryonic control (ECon) or embryonic kynurenine (EKyn; 100 mg kynurenine/day) diet daily. Sleep-wake behavior data were analyzed for 24h on ED 20. **(B)** Wake and total sleep duration in the light phase (Zeitgeber time (ZT) 0-12, white background) or dark phase (ZT 12-24, grey background). **(C)** Relative cage activity in the light phase (ZT 0-12, white background) or dark phase (ZT 12-24, grey background). Data are mean ± SEM with individual data points shown. Significant three-way ANOVA results are displayed in pink annotations below panel B: ++ *p* < 0.01, ++++ *p* < 0.0001. Significant two-way ANOVA results, separated by light phase in panel B, are displayed in red boxed annotations below each panel: ^ *p* < 0.05, ^^^^ *p* < 0.0001. Fisher’s LSD post-hoc: * *p* < 0.05. N = 5-6 dams/group.

### 3.6. EKyn does not elevate inflammation in maternal plasma

To better understand the prenatal phenotypic and biochemical changes induced by EKyn, we collected tissues in a separate cohort of animals. EKyn diet did not induce any significant changes to fetal weight, placental weight, placental efficiency, litter size, or the sex ratio of viable fetuses (**Supplemental Figure 9**). EKyn, compared to ECon, diet did not elevate peripheral inflammatory markers studied in a 27-multiplex assay of maternal plasma (**Figure 8B**, individual means and p-values from t-tests comparing ECon to EKyn are presented in **Supplemental Table 2**). To align the analysis with cytokine markers in the SleepFrag study, data are shown for IFN-γ (**Figure 8C**), IL-1β (**Figure 8D**), IL-6 (**Figure 8E**), and IL-17A (**Figure 8F**).

**Figure 8:**
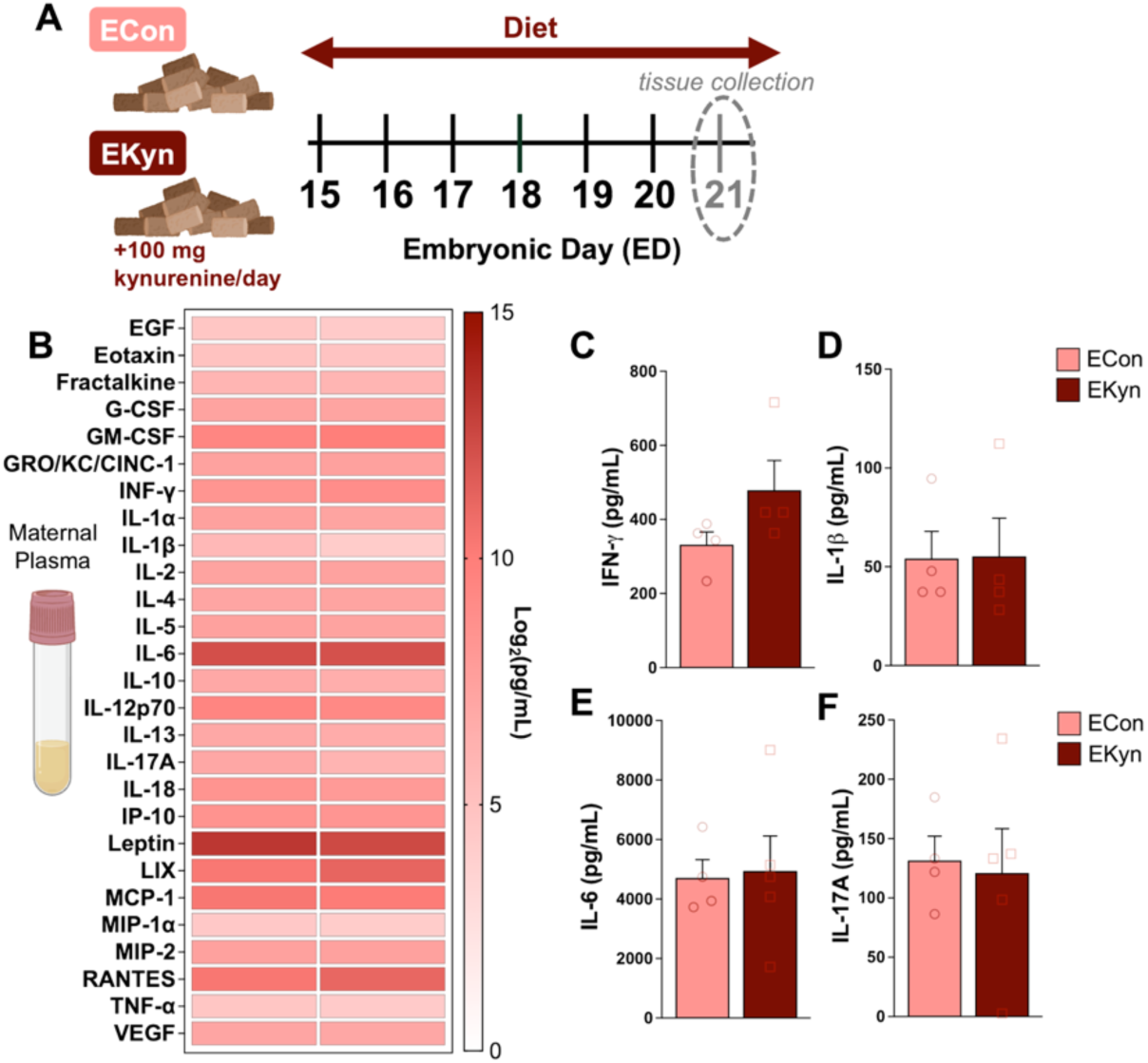
In dams, EKyn diet does not elevate plasma cytokines. **(A)** Schematic of experimental paradigm. From embryonic day (ED) 15 to ED 21, pregnant females received embryonic control (ECon) or embryonic kynurenine (EKyn; 100 mg kynurenine/day) diet daily. Tissues were collected at Zeitgeber time (ZT) 6 on ED 21. Maternal plasma **(B)** 27-plex cytokine discovery panel, **(C)** IFN-γ, **(D)** IL-1β, **(E)** IL-10, and **(F)** IL-17A. Data are mean ± SEM with individual data points shown. No significant results were determined with Mann-Whitney tests. N= 4-5 dams/group.

### 3.7. EKyn elevates KYNA in maternal and fetal tissues

We assessed tryptophan, kynurenine, and KYNA in the maternal plasma. Maternal plasma tryptophan did not change with EKyn diet (**Figure 9A**), yet plasma kynurenine elevated 4.8-fold (**Figure 9B**, *p* < 0.001), plasma QUIN did not differ (**Figure 9C**), and plasma KYNA elevated 2-fold (**Figure 9D**, p < 0.01) and the KYNA/QUIN ratio elevated 2-fold (**Figure 9E**). Kynurenine/tryptophan, QUIN/kynurenine, and KYNA/kynurenine ratios are presented in **Supplemental Figure 10**. Maternal brain kynurenine (**Figure 9F**) and KYNA (**Figure 9G**) levels significantly increased with EKyn compared to ECon in the hypothalamus (kynurenine: +3.7-fold, *p* < 0.01, KYNA: +5.9-fold, *p* < 0.01), frontal cortex (kynurenine: +2.8-fold, *p* < 0.05, KYNA: +2.8-fold, *p* < 0.05) and hippocampus (kynurenine: +2.7-fold, *p* < 0.05, KYNA: +1.8-fold *p* < 0.05).

**Figure 9:**
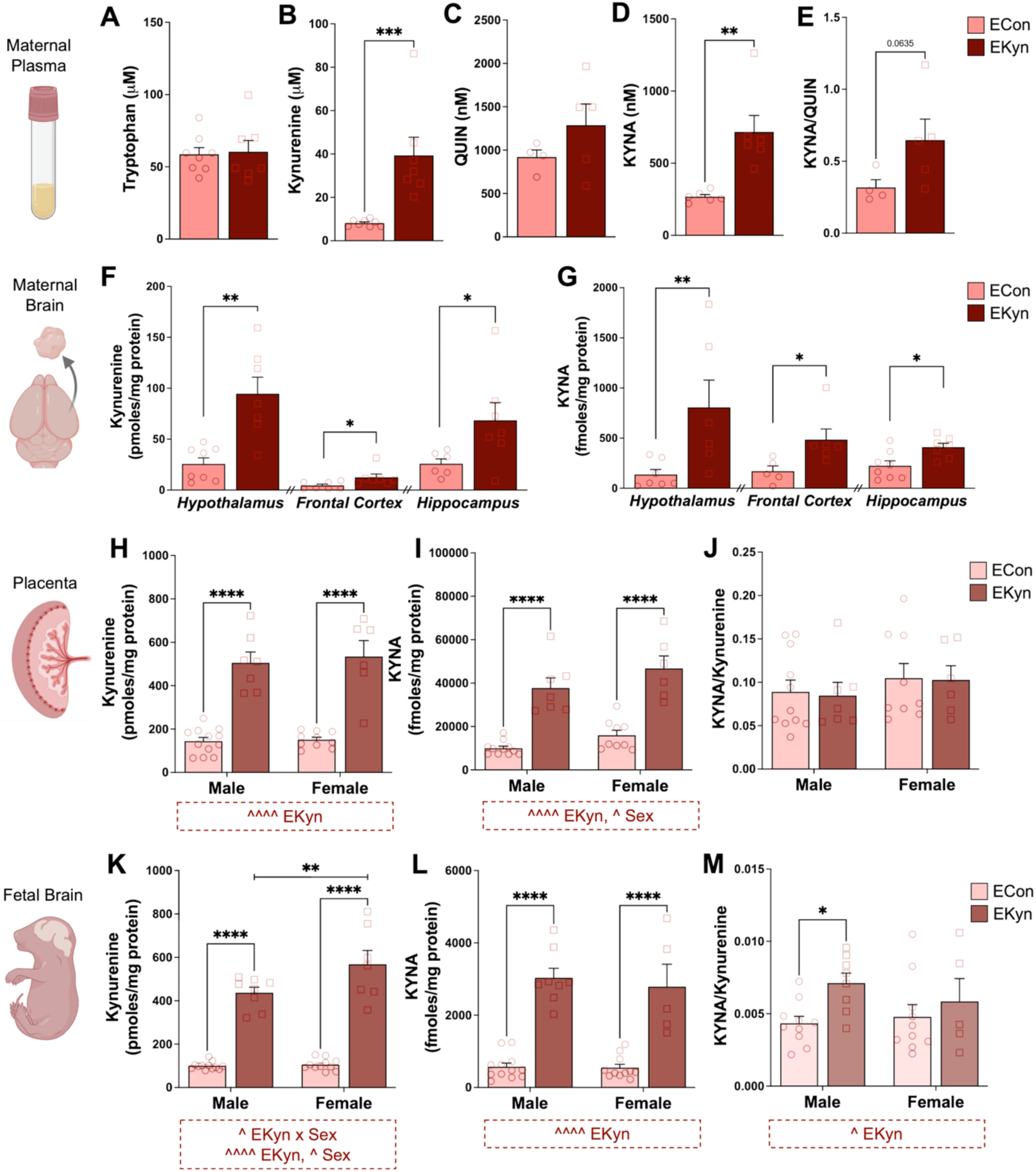
EKyn diet increases tryptophan metabolism toward kynurenine and KYNA in maternal (plasma, brain) and fetal (placenta, brain) tissues. From embryonic day (ED) 15 to ED 21, pregnant females received embryonic control (ECon) or embryonic kynurenine (EKyn; 100 mg kynurenine/day) diet daily. Tissues were collected at Zeitgeber time (ZT) 6 on ED 21. Maternal plasma **(A)** tryptophan, **(B)** kynurenine, **(C)** QUIN, **(D)** KYNA, **(E)** KYNA/QUIN ratio. Maternal brain **(F)** kynurenine **(G)** KYNA. Placental **(H)** kynurenine, **(I)** KYNA, **(J)** KYNA/kynurenine ratio. Fetal brain **(K)** kynurenine, **(L)** KYNA, **(M)** KYNA/kynurenine ratio. Data are mean ± SEM with individual data points shown. Significant two-way ANOVA results are displayed in red boxed annotations below figure panels: ^ *p* < 0.05, ^^^^ *p* < 0.0001. Mann-Whitney test or Fisher’s LSD post-hoc: * *p* < 0.05, ** *p* < 0.01, *** *p* < 0.001, **** *p* < 0.0001. N = 5-10 pups/group, N = 4-8 dams/group.

In the placenta, EKyn did not affect tryptophan levels (**Supplemental Table 3**). EKyn (F_1, 31_ = 110.0, *p* < 0.0001) elevated kynurenine levels 3.5-fold in both males (*p* < 0.0001) and females (*p* < 0.0001) compared to ECon (**Figure 9H**). Of note, placental KYNA levels were impacted by EKyn (F_1, 29_ = 80.50, *p* < 0.0001) and biological sex (F_1, 29_ = 5.274, *p* = 0.0291) (**Figure 9I**). EKyn placentas had higher KYNA in males (+3.8-fold, *p* < 0.0001) and females (+2.9-fold, *p* < 0.0001). Female placentas had overall higher KYNA than male placentas. The KYNA/kynurenine ratio was not changed with EKyn compared to ECon (**Figure 9J**).

Fetal brain kynurenine levels were impacted by an EKyn x sex interaction (F_1, 34_ = 5.647, *p* = 0.0233) and main effects of EKyn (F_1, 34_ = 228.1, *p* < 0.0001) and sex (F_1, 34_ = 6.666, *p* = 0.0143) (**Figure 9K**). EKyn significantly elevated kynurenine levels in the male (+4.4-fold, *p* < 0.0001) and female fetal brain (+5.4-fold, *p* < 0.0001). Female EKyn fetal brain kynurenine levels were significantly higher than those in male EKyn (*p* < 0.01). Fetal brain KYNA was affected only by a main effect of EKyn (F_1, 32_ = 101.8, *p* < 0.0001) (**Figure 9L**). Fetal brain KYNA levels were higher in both male (+5.3-fold, *p* < 0.0001) and female (+5.1-fold, *p* < 0.0001) EKyn brains compared to ECon. Notably, EKyn male fetal brains (*p* < 0.05), but not EKyn female fetal brains, had a higher KYNA/kynurenine ratio than ECon **Figure 9M**).

## 4. Discussion

Our study aimed to characterize the immediate effects of chronic maternal sleep disruption on inflammation and KP metabolism in a sex-dependent manner in utero. To distinguish the effects of maternal sleep disruption from those of elevated KP activity, we employed two independent experimental models during the last week of gestation: 1) maternal SleepFrag, to directly disrupt maternal sleep, and 2) EKyn, to directly enhance KP metabolism. These models were conducted in separate cohorts, allowing us to assess whether distinct maternal and metabolic perturbations converge on similar fetal outcomes, particularly elevated fetal brain KYNA levels. We found that maternal SleepFrag increased maternal plasma inflammatory markers, placental KP metabolism, and fetal brain KYNA, without altering KP metabolism in maternal plasma or brain. These outcomes suggest a unique fetal consequence of KP activation, specifically elevated fetal brain KYNA, following maternal SleepFrag during the last week of gestation. Findings from the EKyn model indicate that increased KP substrate availability alone can drive fetal brain KYNA elevation, with stronger de novo production in males, despite minimal effects on maternal sleep-wake architecture or inflammation. Together, these independent models identify fetal brain KYNA elevation, particularly in males, as a convergent consequence of distinct prenatal insults and highlight the KP as a potential mediator of sex-specific fetal neurodevelopmental vulnerability (de Souza et al., 2026; Gustafsson et al., 2026; Murakami et al., 2023; Notarangelo and Pocivavsek, 2017; van Zundert et al., 2022).

### Consequences of Maternal SleepFrag – Maternal Plasma and Brain

Maternal SleepFrag elevated inflammatory markers in maternal plasma and male placentas at ED 21 following six, but not three, days of sleep disruption. Consistent with our findings, previous studies using maternal sleep deprivation paradigms have observed increased cytokine levels in maternal plasma and placenta (Baratta et al., 2020; Li et al., 2023). Notably, cytokine elevations in the present study emerged only after prolonged SleepFrag exposure, suggesting that sustained sleep disturbance may be necessary to trigger inflammatory changes during pregnancy (Garbarino et al., 2021). Specifically, SleepFrag increased IFN-γ, IL-17A, and IL-6 in maternal plasma and selectively elevated IL-6 in male placentas. Because even modest increases in pro-inflammatory cytokines have been associated with altered fetal neurodevelopment, these findings suggest that maternal sleep disruption may create an inflammatory environment with long-term consequences for offspring (Choi et al., 2016; Han et al., 2021; Meyer, 2014; Otero and Antonson, 2022; Wu et al., 2017).

An elevated prenatal inflammatory environment has been linked to increased KP metabolism, including from preclinical models such as prenatal lipopolysaccharide (LPS) exposure or restraint stress (Abad et al., 2024; Abad et al., 2025; Hasegawa et al., 2024; Notarangelo and Schwarcz, 2016, 2021; Petrova et al., 2024; Santana-Coelho et al., 2026; Sha et al., 2024). Mechanistically, pro-inflammatory cytokine signaling, particularly from IFN-γ, can enhance IDO1 transcription, a rate-limiting enzyme that converts tryptophan to kynurenine, thereby increasing kynurenine availability and downstream KP metabolites in peripheral tissues and the brain (Stone et al., 2022; Stone and Williams, 2023; Wang et al., 2022). Although SleepFrag induced a maternal inflammatory response, KP metabolites were not increased in maternal plasma or brain. This finding is consistent with previous work from our laboratory, where acute sleep deprivation did not increase KP metabolites in female rats or pregnant dams (Baratta et al., 2018a; Baratta et al., 2020). Although we suggested that acute sleep loss in females may engage corticosterone-related mechanisms, plasma corticosterone levels remained unchanged in the present study. This aligns with prolonged sleep-restriction paradigms in which hypothalamic-pituitary-adrenal (HPA)-axis activation becomes blunted, a response that may, during pregnancy, serve as a protective adaptation for the developing fetus (Pardo et al., 2016; Sze and Brunton, 2024; Wright et al., 2023b). Together, these findings suggest that chronic SleepFrag-induced inflammation is not sufficient to broadly activate KP metabolism within maternal compartments and further support a model of relative female resilience to sleep-disturbance-induced KP alterations, even during pregnancy.

### Consequences of Maternal SleepFrag - Placenta

In contrast to the maternal compartment, SleepFrag elevated KP metabolism in fetal tissues, as reflected by increased tryptophan and kynurenine levels in the placenta and fetal brain. Because the placenta is highly vascularized and these metabolites can cross the placental barrier, residual maternal blood could potentially influence placental metabolite measurements (Goeden et al., 2017; Notarangelo and Pocivavsek, 2017). However, this is unlikely to fully explain the present findings, as maternal plasma tryptophan and kynurenine levels were unchanged following SleepFrag. Further, an increased kynurenine/tryptophan ratio, a proximal marker of IDO1/TDO activity (Badawy and Guillemin, 2019), was observed specifically in the placenta. Together, these findings suggest that SleepFrag activates KP metabolism in a fetal-compartment-specific manner, likely beginning at the placenta, with subsequent transfer of tryptophan and kynurenine to the fetal brain, where KYNA can be produced *de novo* (Goeden et al., 2017; Notarangelo et al., 2019).

The placenta is a critical mediator of the maternal-fetal interface and plays a central role in regulating fetal physiology (Kramer et al., 2023). Prenatal insults, including maternal sleep loss and inflammation, can impair placental function and disrupt fetal neurodevelopment (Bronson and Bale, 2016; Goeden et al., 2016; Woods et al., 2023). Thus, SleepFrag-induced activation of placental KP metabolism may represent one mechanism by which maternal sleep disruption alters the fetal environment and contributes to neurodevelopmental vulnerability (Baratta et al., 2020; Karahoda et al., 2026; Wright et al., 2023b).

Elevated placental tryptophan following SleepFrag, despite unchanged maternal plasma tryptophan, is notable given that tryptophan is an essential amino acid and a precursor for multiple bioactive pathways. One possibility is that SleepFrag alters maternal or placental tryptophan handling through changes in gut-derived tryptophan metabolism, placental amino acid transport, or serotonin/melatonin synthesis, all of which are sensitive to sleep disturbance and inflammation (Agus et al., 2018; Galley et al., 2021; Goeden et al., 2016; Pires et al., 2021; Wang and Li, 2021). Alternatively, increased placental tryptophan availability may augment serotonin synthesis (Goeden et al., 2016). This is particularly relevant because the placenta is the primary source of fetal serotonin during late rodent gestation and increased placental serotonin metabolism during this developmental period can interfere with essential neurogenic processes in the fetal brain (Bonnin and Levitt, 2011; Goeden et al., 2016). While future studies should directly assess these alternative considerations, a significant negative correlation (data not shown) between maternal plasma and placental tryptophan levels suggests enhanced maternal-to-placental transfer following SleepFrag. Together with the increase in placental kynurenine and fetal brain KP metabolites described above, these findings support the interpretation that SleepFrag preferentially alters tryptophan metabolism within the fetal compartment

### Consequences of Maternal SleepFrag – Fetal Brain

Although SleepFrag elevated KP metabolites across fetal tissues, KYNA elevation was confined to the fetal brain, suggesting tissue-specific regulation of KP metabolism during prenatal SleepFrag. Activation of the KP during inflammation may serve a protective function by facilitating negative feedback on inflammatory signaling (Mándi and Vécsei, 2012; Zhai et al., 2020). Moreover, fetal KP enzymes and metabolites, including KYNA levels, are naturally elevated during prenatal development, which is hypothesized to support critical developmental processes, such as preventing maternal T cell activation and fetal immune rejection and protecting from hypoxia during birth (Notarangelo and Pocivavsek, 2017). Nevertheless, preclinical studies have linked excessive or prolonged elevation of fetal brain KYNA to lasting neurochemical and behavioral deficits, including sleep and cognitive disturbances, in young adult offspring (Beggiato et al., 2025; Hahn et al., 2018; Milosavljevic et al., 2025; Milosavljevic et al., 2023; Notarangelo and Pocivavsek, 2017; Pershing et al., 2015; Pershing et al., 2016; Pocivavsek et al., 2014; Pocivavsek et al., 2012; Rentschler et al., 2021; Wright et al., 2021). To the best of our knowledge, the present findings are the first to identify elevated fetal brain KYNA as a consequence of chronic maternal SleepFrag.

The precise mechanism through which elevated fetal brain KYNA alters neurodevelopment remains to be elucidated; however, the leading hypothesis involves antagonism of α7nACh and NMDA receptors (Bagasrawala et al., 2016; Beggiato et al., 2025; Notarangelo and Pocivavsek, 2017; Schwarcz et al., 2012), both of which are integral to neurodevelopmental regulation (Abreu-Villaça et al., 2011; Van Dongen, 2009; Wu et al., 2015). Although less comprehensively understood, particularly in the context of pregnancy, activation of GPR35 or AhR could also contribute (DiNatale et al., 2010; Moroni et al., 2012; Pocivavsek et al., 2016; Wang et al., 2006).

### Considerations for Fetal Sex

A major strength of the present study is the ability to sex-match placental and fetal tissues, enabling us to evaluate sex-specific effects of prenatal maternal sleep disruption and elevated kynurenine exposure in utero. This approach revealed increased male vulnerability across inflammatory and KP outcomes, which aligns with numerous preclinical models of prenatal insults relevant to neurodevelopmental disorders (Bale, 2016; Hall et al., 2023). The placenta is an important site of sexual dimorphism, and male and female placentas differ in their translational, metabolic, and inflammatory responses to prenatal insults (Baines and West, 2023; Bale, 2016; Bordt et al., 2021; Paparini et al., 2024). Female placentas tend to be more immune-tolerant and adaptable than male placentas (Baines and West, 2023; Bale, 2016), which aligns with our finding that SleepFrag elevated IL-6 in male placentas only. However, both male and female placentas exhibited increased tryptophan, kynurenine, and kynurenine/tryptophan ratios following SleepFrag, suggesting that mechanisms such as undetectable or unmeasured cytokines, reactive oxygen species or other secondary metabolites, and corticosterone may contribute to placental IDO1/TDO activation, particularly in females (Badawy, 2017; Blanco Ayala et al., 2015).

Sex differences were also evident in the fetal brain. SleepFrag elevated fetal brain kynurenine in both males and females, yet elevated KYNA to a greater extent in males. EKyn elevated fetal brain kynurenine in both sexes, whereas the KYNA/kynurenine ratio was increased only in males, suggesting heightened KAT activity or KYNA synthesis in male fetal brains. To our knowledge, this is the first study to identify sex-specific regulation of KP metabolism in utero, constituting the need for future studies to investigate whether differential activity of distinct KAT isoforms or alternative routes of KYNA production, such as non-enzymatic oxidation, contribute to sex differences in fetal brain KYNA production (Al-Gubory and Garrel, 2016; Badawy, 2017; Blanco Ayala et al., 2015). Future work with maternal SleepFrag and EKyn will also consider whether males and females differentially engage the alternative KMO branch of the KP that produces 3-hydroxykynurenine (3-HK) and QUIN, metabolites that can promote oxidative stress and neurotoxicity (Foster et al., 1984; Guillemin, 2012; Schwarcz et al., 2012). Sex-biased utilization of distinct KP branches could contribute to divergent neurodevelopmental outcomes between males and females. Together, these findings highlight fetal and placental sex determination as a critical approach for uncovering biological mechanisms that may be masked in mixed-sex analyses, including prior studies of prenatal KP dynamics (Baratta et al., 2020; Goeden et al., 2017; Notarangelo et al., 2019; Notarangelo and Pocivavsek, 2017; Notarangelo and Schwarcz, 2021; Silvano et al., 2021). Given that adult male EKyn offspring exhibit more pronounced behavioral and neurochemical impairments than females (Buck et al., 2020; Rentschler et al., 2021; Wright et al., 2021), the current sex-specific findings in SleepFrag and EKyn models provide an important foundation for investigating the mechanisms underlying sexual divergence following prenatal insults.

### Consequences of EKyn – Maternal Inflammation and Sleep-wake Behaviors

Finally, we evaluated inflammation and maternal sleep-wake behavior in EKyn dams to determine whether elevated kynurenine availability disrupts maternal physiology during pregnancy. The EKyn diet did not affect inflammatory markers in maternal plasma, despite emerging evidence that KP metabolites can modulate inflammation under certain conditions (Silvano et al., 2021; Stone et al., 2022; Stone and Williams, 2023). EKyn also did not produce major disruption in sleep-wake architecture observed with SleepFrag as we saw no change in sleep-wake durations in the light phase, when rodents accumulate the most sleep. EKyn did increase activity and wakefulness during the dark phase. This relatively modest arousal phenotype was unexpected given prior work from our laboratory demonstrating a causal role for KYNA in sleep regulation, in which increased brain KYNA levels directly disrupt light phase sleep-wake architecture, and reduced levels enhance sleep behaviors (Milosavljevic et al., 2023; Pocivavsek et al., 2017; Rentschler et al., 2024). Although brain KYNA was elevated in EKyn dams, light phase sleep-wake architecture remained intact, suggesting that pregnancy may alter the relationship between KP activation and sleep regulation. Pregnancy-associated changes in steroid hormones, immune regulation, and neurotransmission may influence sleep regulation and should be considered as potential mediators of KP effects during pregnancy (Chechko et al., 2022; Servin-Barthet et al., 2023; Solano and Arck, 2019). Indeed, ongoing work from our laboratory indicates that sleep dynamics and KP sensitivity differ substantially between pregnant and non-pregnant females, underscoring the need for future studies focused specifically on pregnancy-dependent regulation of KP-sleep interactions (Wright et al., 2023a). Importantly, the absence of maternal inflammation and major sleep disruption in EKyn dams strengthens the interpretation that elevated fetal brain KYNA levels, rather than maternal inflammatory or sleep-related changes, contribute to the later behavioral and neurochemical deficits reported in EKyn offspring (Beggiato et al., 2025; Buck et al., 2020; Hahn et al., 2018; Pershing et al., 2015; Pershing et al., 2016; Pocivavsek et al., 2014; Pocivavsek et al., 2012; Rentschler et al., 2021; Wright et al., 2021).

### Conclusions

Together, our studies of SleepFrag and EKyn identify elevated fetal brain KYNA as a convergent neurochemical outcome through which maternal sleep disruption, inflammation, and/or KP activation may influence sex-specific fetal brain development (**Figure 10**). These results also support the use of our EKyn model as a translation tool to isolate the neurodevelopmental consequences of increased prenatal KP metabolism. Clinically, maternal sleep disruption is common during pregnancy and is associated with adverse maternal and offspring outcomes, yet the fetal biochemical mechanisms linking disrupted maternal sleep to neurodevelopmental risk remain poorly understood (Carroll et al., 2019; Satyapal et al., 2026; Wright et al., 2023b). Our findings suggest that fetal brain KP metabolism, and KYNA in particular, may represent a mechanistic link between prenatal environmental insults and altered neurodevelopmental trajectories. Ultimately, protecting maternal sleep and maintaining appropriate regulation of fetal brain KYNA levels may be important considerations for promoting fetal health throughout pregnancy.

**Figure 10:**
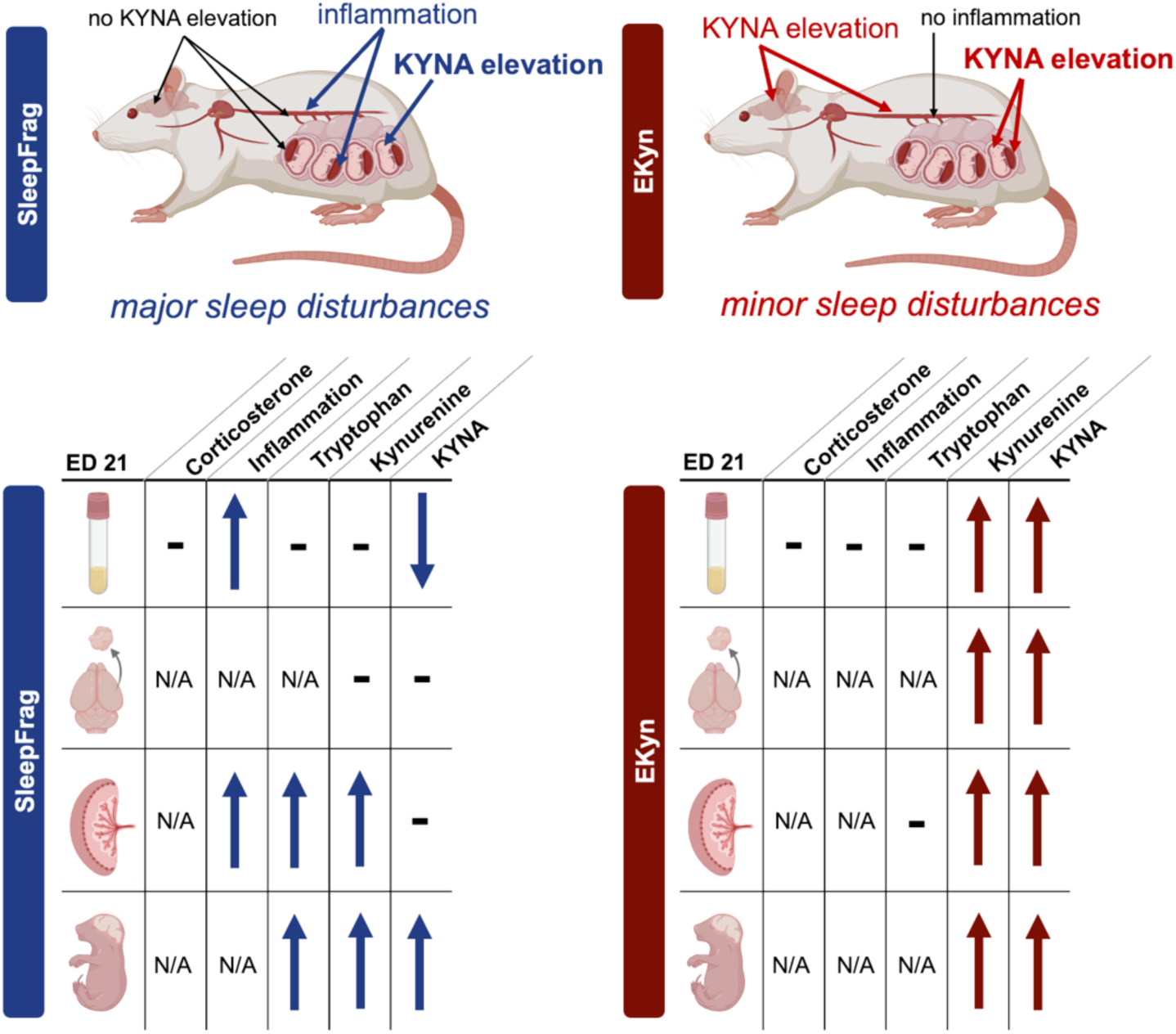
Maternal sleep disruption (SleepFrag) and kynurenine diet (EKyn) differentially alter prenatal physiology but converge on increased fetal brain KYNA. Maternal sleep SleepFrag markedly disrupts maternal sleep, increases peripheral and placental inflammation, and elevates fetal brain KYNA. In contrast, EKyn elevates fetal brain KYNA in the absence of major maternal sleep disruption or inflammation. Across both models, effects are more pronounced in males than in females. Collectively, these complementary models dissociate the contributions of maternal sleep disruption and kynurenine pathway (KP) activation while identifying fetal brain KYNA elevation as a common outcome important to neurodevelopment.

## Acknowledgments

The authors would like to acknowledge Matthew Efird, Charles A. Grant, Gannett L. Loftus, Maria V. Piroli, Sam C. Walther, and Corbin E. Witt for their technical contributions to this work. Some figure components were generated in BioRender.

## Author Contributions

**CJW:** Conceptualization, investigation, methodology, formal analysis, visualization, writing – original draft, writing – review & editing, funding acquisition. **JHC:** methodology, investigation, visualization, writing – original draft, writing – review & editing. **SM:** validation, writing – review & editing. **HV:** software, writing – review & editing, funding acquisition. **NF:** methodology, investigation, visualization, writing – original draft, writing – review & editing, project administration, supervision, resources. **AP:** Conceptualization, investigation, methodology, formal analysis, visualization, writing – original draft, writing – review and editing, funding acquisition, project administration, supervision, resources.

---

**Supplemental Figure 1:**
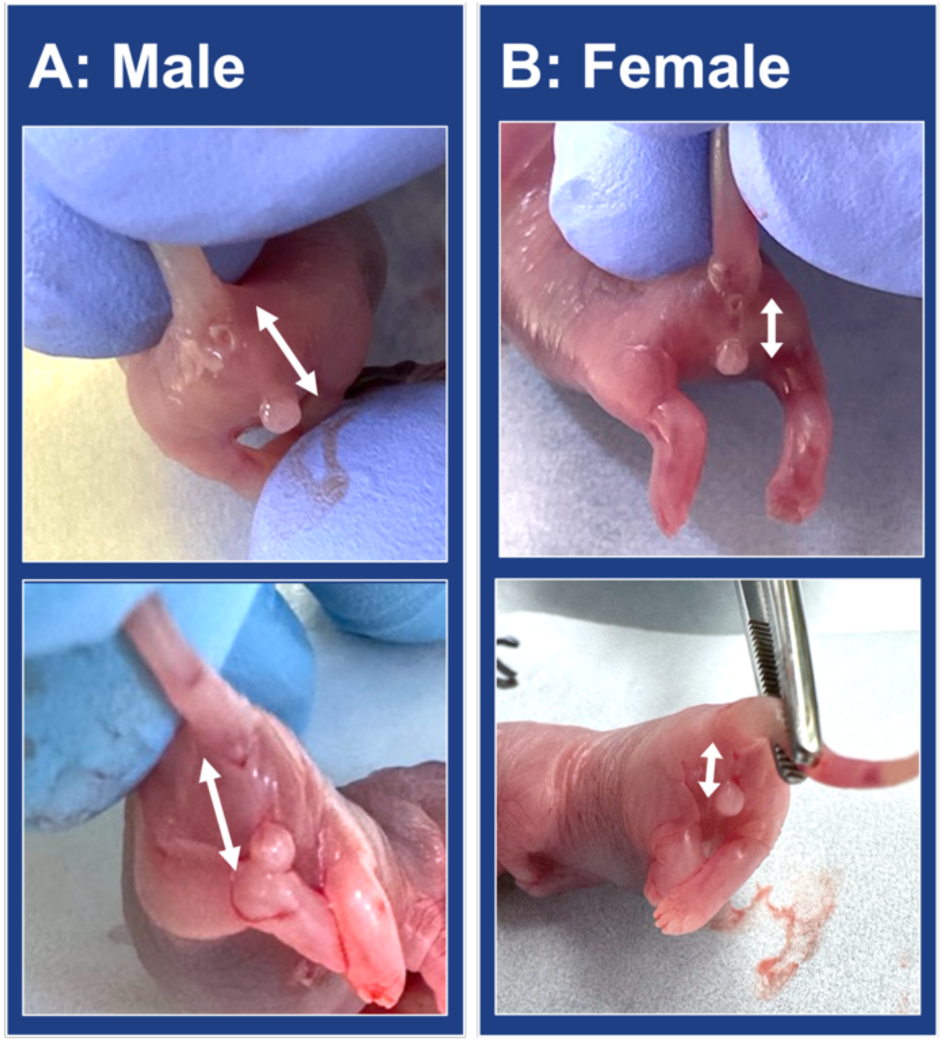
Visual determination of fetal sex at embryonic day (ED) 21. **(A)** Male fetuses are identified by a longer anogenital distance, defined as the distance between the anus and external genitalia. **(B)** Female fetuses are identified by a shorter anogenital distance. White arrows indicated the anogenital distance.

**Supplemental Figure 2:**
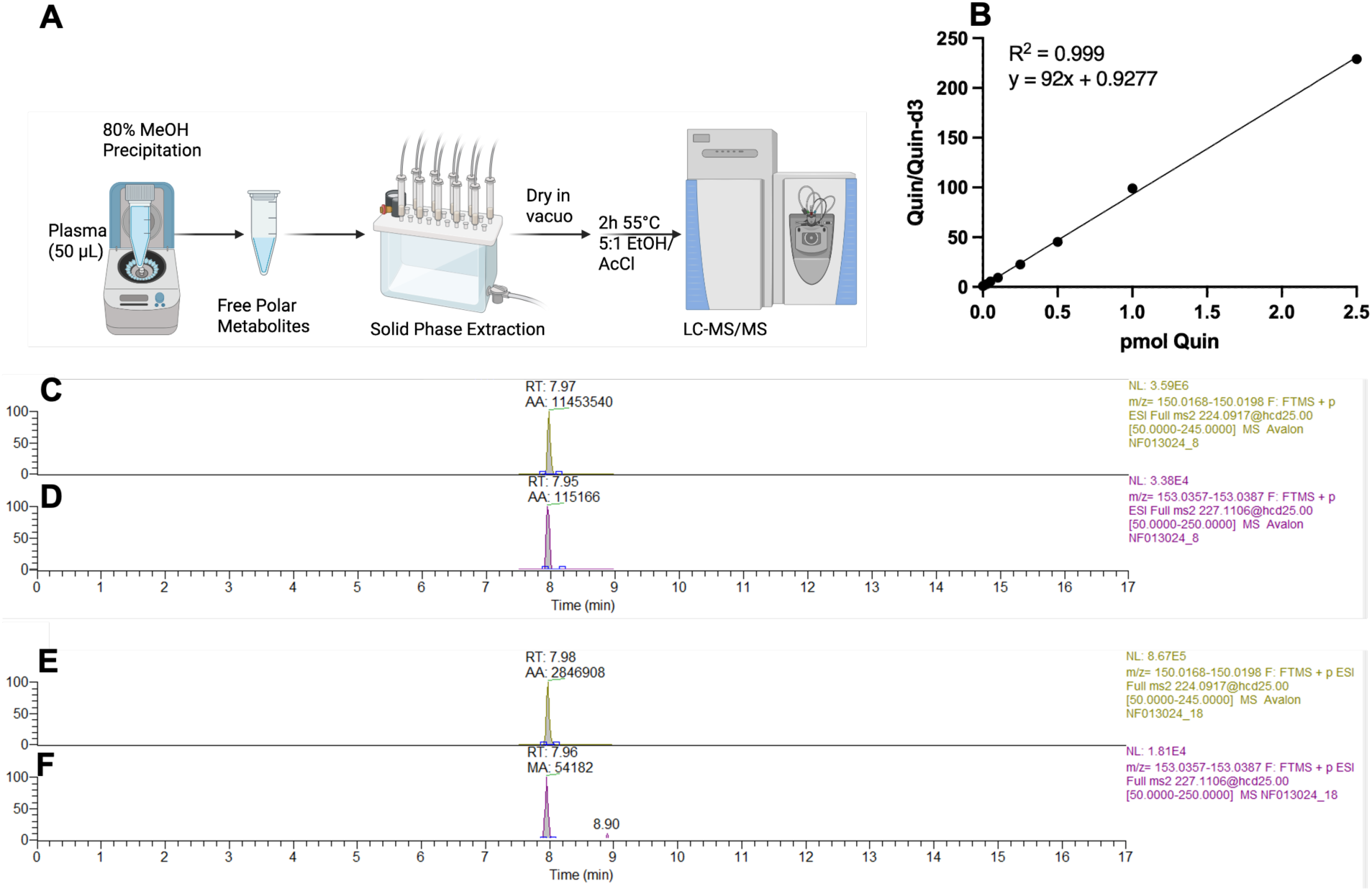
Ethyl esterification of quinolinic acid (QUIN) for LC-MS/MS quantification. **(A)** Schematic of the sample processing workflow. **(B)** Standard curve for QUIN from 5 fmol to 2.5 pmol. Selected ion chromatograms for ethyl esterified **(C)** QUIN and **(D)** QUIN-d3 internal standard from the 1 pmol standard. Corresponding selected ion chromatograms for ethyl esterified **(E)** QUIN and **(F)** QUIN-d3 from a biological sample.

**Supplemental Table 1:** Control vs SleepFrag cytokines.

| Supplemental Table 1: Control vs SleepFrag cytokines |  |  |  |  |
| --- | --- | --- | --- | --- |
| Metabolite | ED 18 |  | ED 21 |  |
|  | Significance | p-value | Significance | p-value |
| EGF | ns | 0.1143 | ns | 0.0952 |
| Eotaxin | ns | 0.7619 | ** | 0.0095 |
| Fractalkine | ns | 0.9143 | ** | 0.0095 |
| G-CSF | ns | 0.2571 | ** | 0.0095 |
| GM-CSF | ns | 0.181 | ** | 0.0095 |
| GRO | ns | 0.4762 | * | 0.0238 |
| INF-γ | ns | 0.1143 | * | 0.0238 |
| IL-1α | ns | 0.1905 | ns | 0.6095 |
| IL-1β | ns | 0.3524 | ns | 0.9048 |
| IL-2 | ns | 0.1952 | * | 0.019 |
| IL-4 | ns | 0.4762 | ** | 0.0048 |
| IL-5 | ns | 0.6095 | ** | 0.0095 |
| IL-6 | ns | 0.1714 | ** | 0.0095 |
| IL-10 | ns | 0.9571 |  |  |
| IL-12p70 | ns | 0.2571 | ** | 0.0095 |
| IL-13 | ns | 0.5762 | ** | 0.0095 |
| IL-17A | ns | 0.1524 | ** | 0.0095 |
| IL-18 | ns | 0.2857 | ** | 0.0095 |
| IP-10 | ns | >0.9999 | * | 0.019 |
| leptin | ns | 0.1143 | ** | 0.0095 |
| LIX | ns | 0.1714 | ns | 0.7619 |
| MCP-1 | ns | >0.9999 |  |  |
| MIP-1α | ns | >0.9999 | ** | 0.0095 |
| MIP-2 | ns | 0.2333 | ** | 0.0095 |
| RANTES | ns | 0.7619 | ns | >0.9999 |
| TNF-α | ns | 0.3238 | | |
| VEGF | ns | 0.7619 | ns | 0.1143 |

**Supplemental Figure 3:**
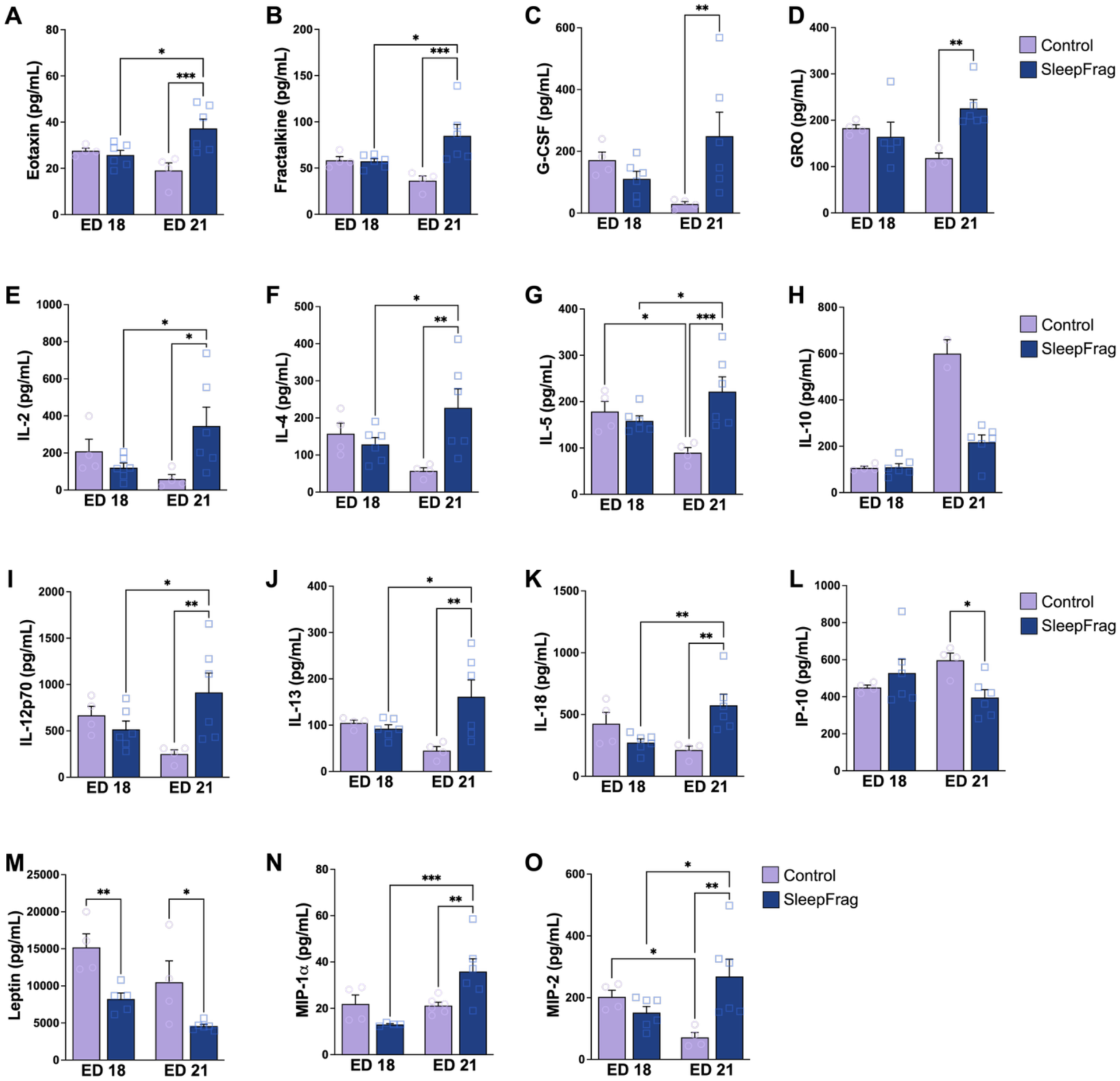
In dams, SleepFrag increases inflammatory markers in maternal plasma during pregnancy. **(A)** Eotaxin **(B)** Fractalkine **(C)** granulocyte colony-stimulating factor (G-CSF) **(D)** growth regulated oncogene (GRO) **(E)** interleukin (IL) 2 **(F)** IL-4 **(G)** IL-5 **(H)** IL-10 **(I)** IL-12p70 **(J)** IL-13 **(K)** IL-18 **(L)** interferon gamma-induced protein (IP)-10 **(M)** Leptin **(N)** macrophage inflammatory protein 1 alpha (MIP1a) **(O)** MIP-2. All data are mean ± SEM with individual data points shown. Two-way ANOVA was conducted, and Fisher’s LSD post-hoc test results are shown on the graphs: * *p* < 0.05, ** *p* < 0.01, *** *p* < 0.001. N = 3-6 dams/group.

**Supplemental Figure 4:**
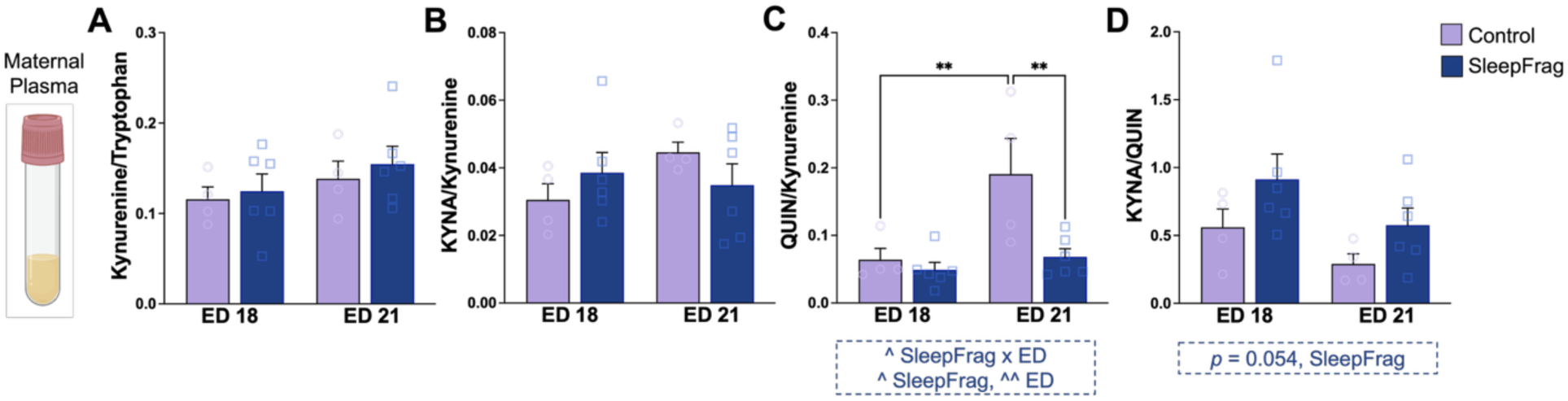
In dams, SleepFrag shifts kynurenine pathway metabolism in plasma towards enhanced KYNA formation. Maternal plasma **(A)** Kynurenine/tryptophan ratio, **(B)** KYNA/kynurenine ratio, **(C)** QUIN/kynurenine ratio, **(D)** KYNA/QUIN ratio. All data are mean ± SEM with individual data points shown. Significant two-way ANOVA results are displayed in blue boxed annotations below each figure panel: ^ *p* < 0.05. Fisher’s LSD post-hoc: ** *p* < 0.01. N = 4-6 dams/group.

**Supplemental Figure 5:**
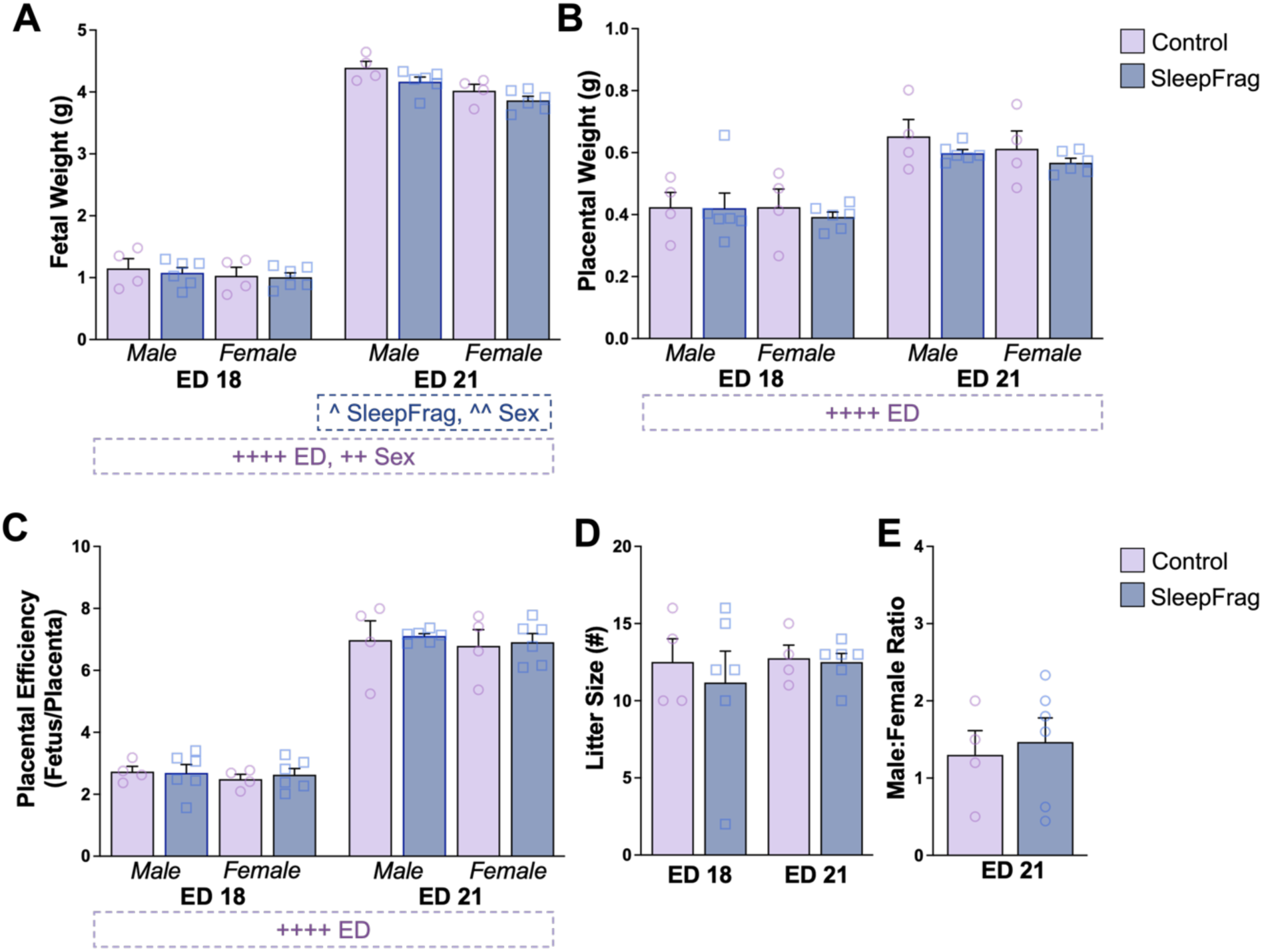
Maternal SleepFrag reduces fetal body weight at embryonic day (ED) 21. **(A)** Fetal body weight **(B)** Placenta weight **(C)** Placental efficiency (fetal body weight/placenta weight) **(D)** Litter size **(E)** Male/female ratio at ED 21. This ratio was only calculated at ED 21 because sex was not determined in all ED 18 fetuses. Data are mean ± SEM with individual data points shown. These individual data points represent litter averages. Significant three-way ANOVA results are displayed in purple annotations below each figure panel: ++ *p* < 0.01, ++++ *p* < 0.0001. Significant two-way ANOVA results, separated by ED in Panel A, are displayed in blue boxed annotations below each figure panel: ^ *p* < 0.05, ^^ *p* < 0.01. No significant results were determined with Fisher’s LSD. N = 4-6 litters/group.

**Supplemental Figure 6.**
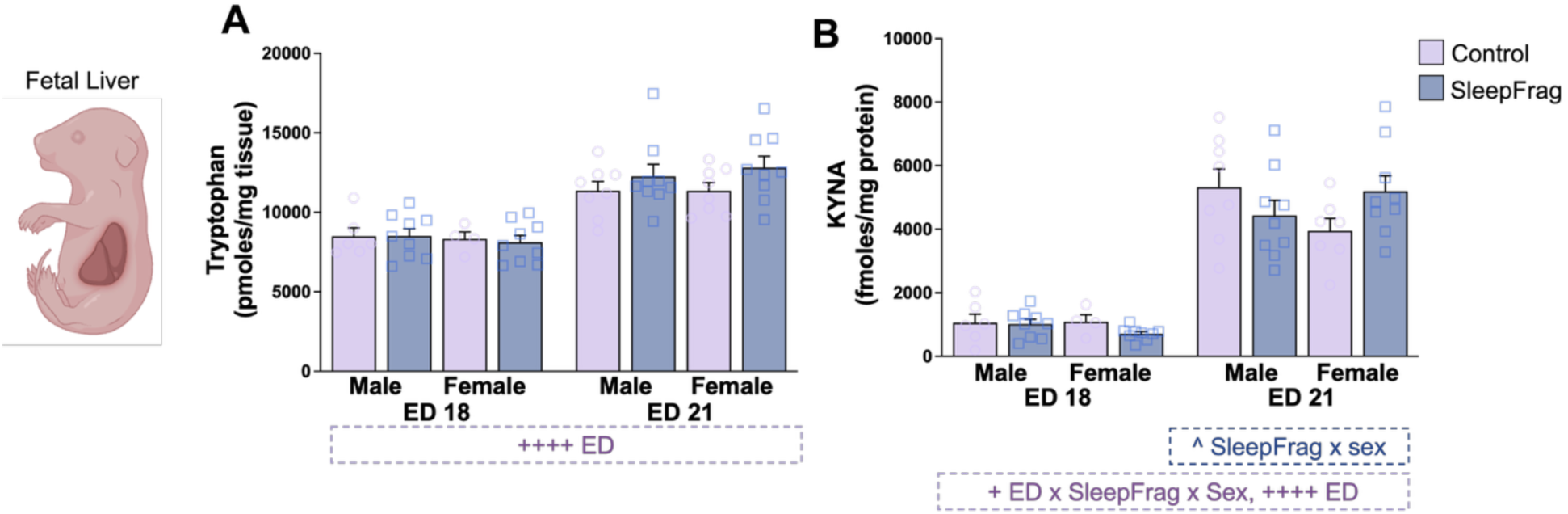
: Maternal SleepFrag elevates tryptophan but not KYNA in the fetal liver at embryonic day (ED) 21. Fetal liver **(A)** tryptophan **(B)** KYNA. Data are mean ± SEM with individual data points shown. Significant three-way ANOVA results are displayed in purple annotations below each figure panel: + *p* < 0.05, ++++ *p* < 0.0001. Significant two-way ANOVA results, separated by ED, are displayed in blue boxed annotations below each figure panel: ^ *p* < 0.05. Fisher’s LSD post-hoc. N = 4-9 pups/group, N = 4-6 dams/group.

**Supplemental Figure 7:**
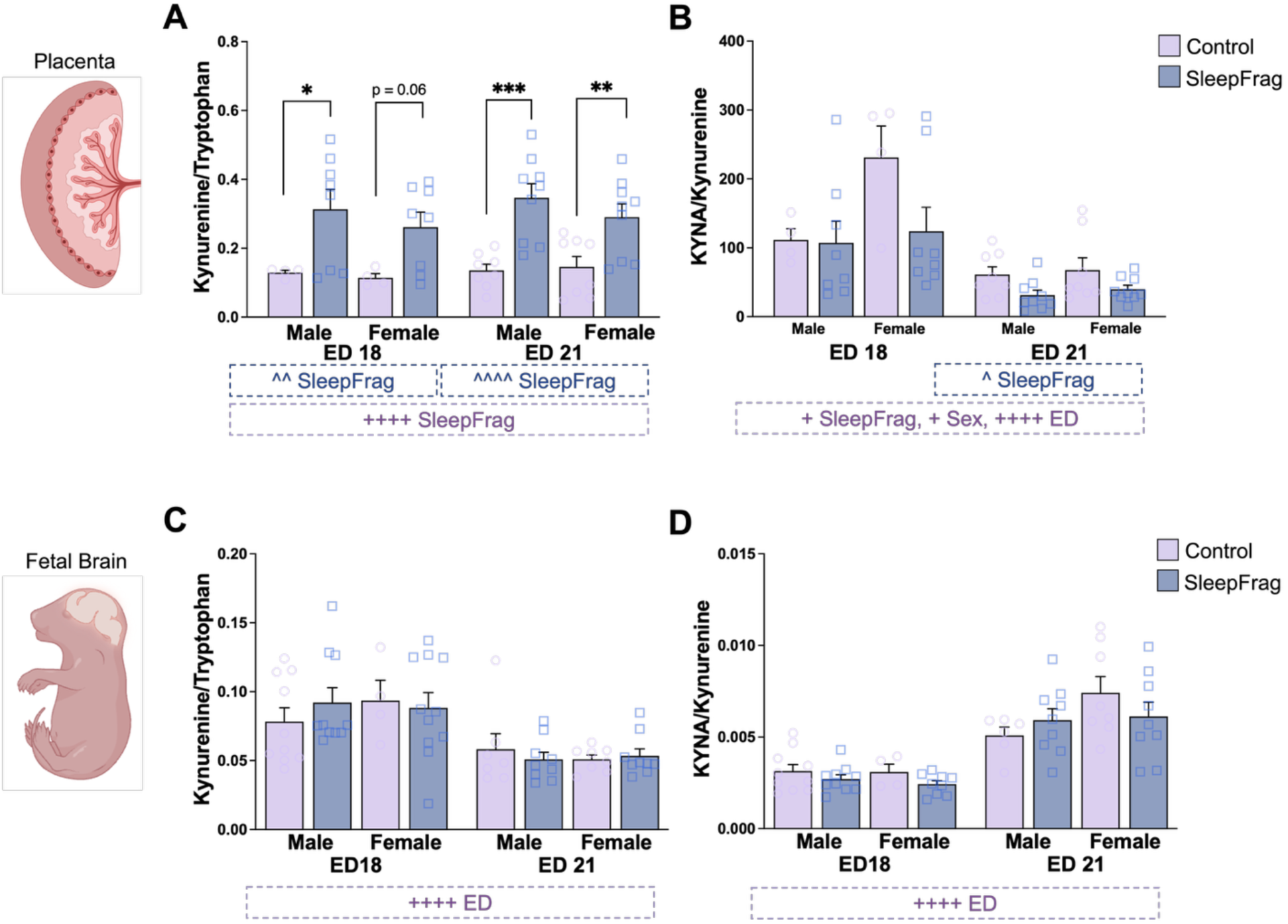
Maternal SleepFrag elevates placental kynurenine pathway metabolism in both sexes but does not affect fetal brain metabolism. Placental **(A)** Kynurenine/tryptophan ratio and **(B)** KYNA/kynurenine ratio. Fetal brain **(C)** Kynurenine/tryptophan ratio and **(D)** KYNA/kynurenine ratio. Data are mean ± SEM with individual data points shown. Significant three-way ANOVA results are displayed in purple annotations below each figure panel: + *p* < 0.05, ++++ *p* < 0.0001. Significant two-way ANOVA results are displayed in blue boxed annotations below each figure panel: ^ *p* < 0.05, ^^ *p* < 0.01. Fisher’s LSD post-hoc: * *p* < 0.05, ** *p* < 0.01 *** *p* < 0.001. N = 4-10 pups/group, N = 4-6 dams/group.

**Supplemental Figure 8:**
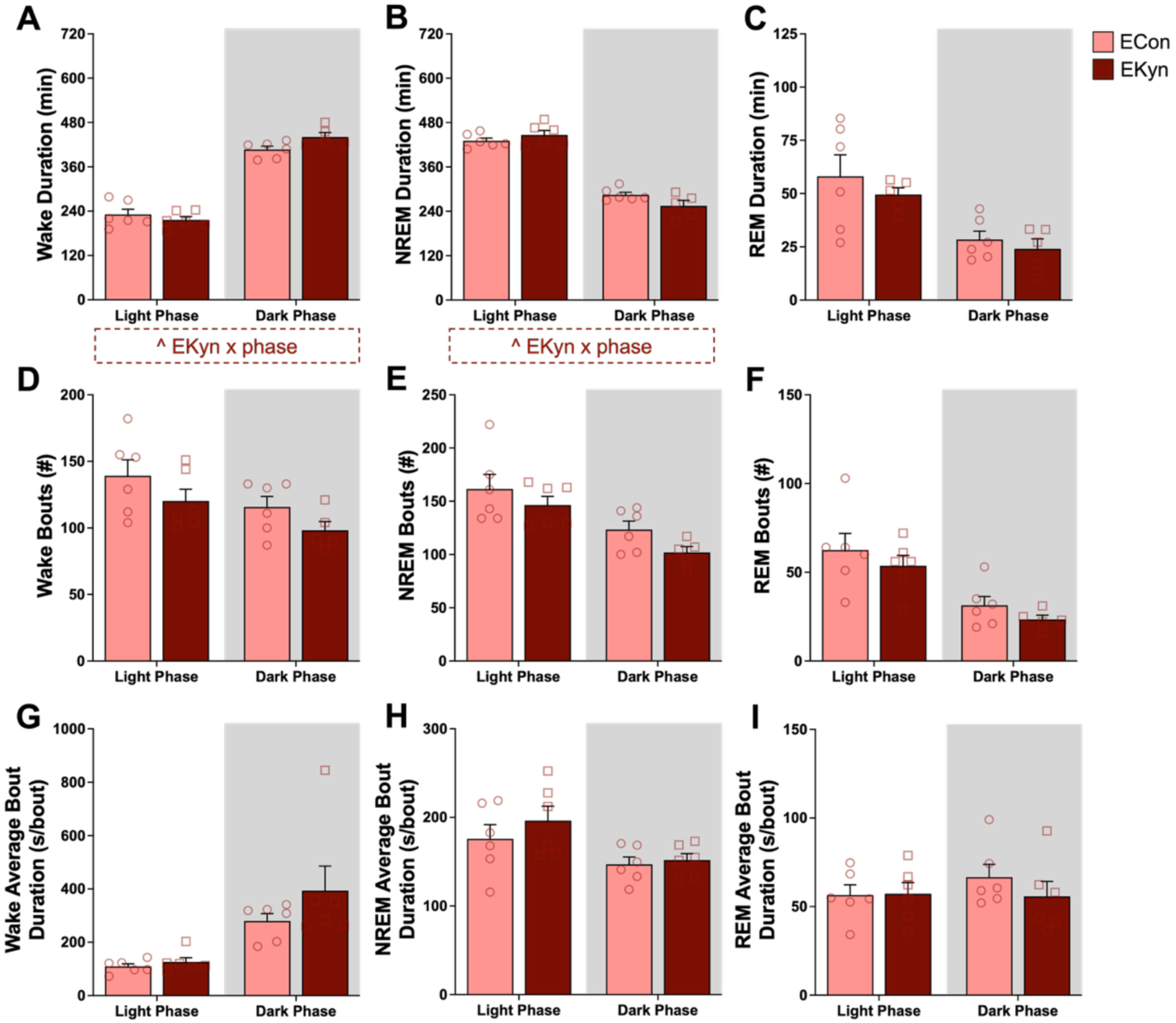
EKyn diet does not significantly alter sleep-wake architecture in dams. Sleep-wake data on embryonic day (ED) 20 were classified as wake, non-rapid eye movement sleep (NREM), or rapid eye movement sleep (REM). Light phase corresponds to white background (zeitgeber time (ZT) 0 – ZT 12) and dark phase corresponds to grey background (ZT 12- ZT 24). **(A)** Wake duration **(B)** NREM duration **(C)** REM duration **(D)** wake bout number **(E)** NREM bout number **(F)** REM bout number **(G)** wake average bout duration **(H)** NREM average bout duration. **(I)** REM average bout duration. Data are mean ± SEM with individual data points shown. Significant two-way RM ANOVA results are displayed in red boxed annotations below each figure panel: ^ *p* < 0.05. No significant results were determined with Fisher’s LSD post hoc. N = 5-6 dams/group.

**Supplemental Figure 9:**
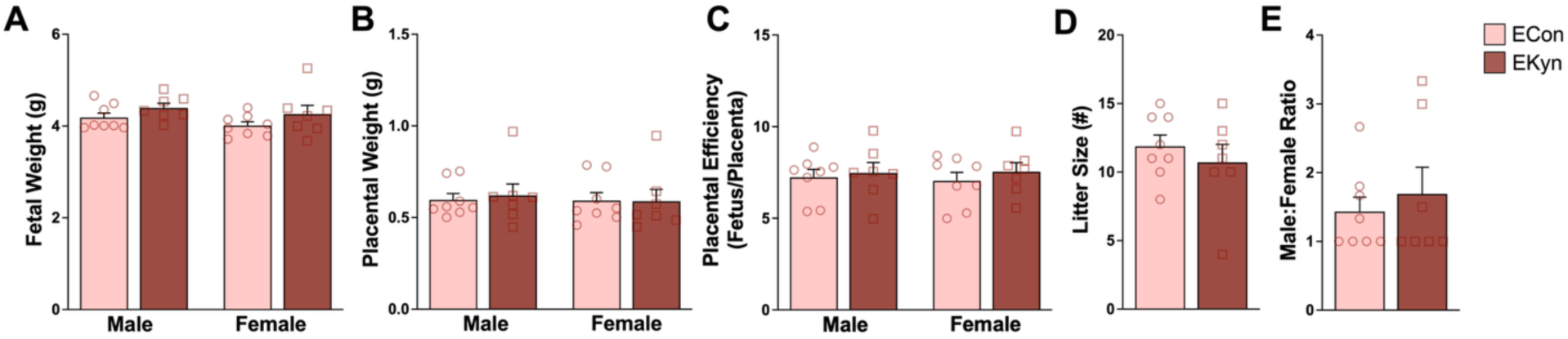
EKyn diet does not produce gross developmental alterations. **(A)** Fetal body weight, **(B)** placental weight, **(C)** placental efficiency (fetal body weight/placenta weight), **(D)** litter size, and **(E)** Male/female ratio. Data are mean ± SEM with individual data points shown. These individual data points represent litter averages. No significant differences were determined with two-way ANOVA or Mann-Whitney tests. N = 7-8 litters/group.

**Supplemental Table 2:** ECon vs EKyn cytokines (pg/mL)

| *Metabolite* | *ECon*<br>*(mean ± SEM)* | *EKyn*<br>*(mean ± SEM)* | *ECon vs EKyn*<br>*Significance* | *p-value* |
| --- | --- | --- | --- | --- |
| EGF | 51.54 ± 34.83 | 43.22 ± 23.56 | ns | 0.9048 |
| Eotaxin | 30.58 ± 1.604 | 30.4 ± 5.094 | ns | 0.9048 |
| Fractalkine | 59.88 ± 3.491 | 64.22 ± 11.57 | ns | 0.9048 |
| G-CSF | 161.9 ± 27.41 | 196.6 ± 60.68 | ns | 0.9048 |
| GM-CSF | 664.9 ± 57.22 | 995.8 ± 183.6 | ns | 0.1429 |
| GRO | 167 ± 10.84 | 181.8 ± 31.87 | ns | 0.5556 |
| INF-γ | 332.1 ± 34.08 | 479.3 ± 79.94 | ns | 0.0857 |
| IL-1α | 176.4 ± 17.15 | 199.8 ± 61.67 | ns | >0.9999 |
| IL-1β | 54.23 ± 13.68 | 44.32 ± 18.52 | ns | 0.5476 |
| IL-2 | 163.5 ± 24.82 | 221.8 ± 84.86 | ns | 0.9048 |
| IL-4 | 138 ± 14.29 | 172.1 ± 42.52 | ns | 0.7302 |
| IL-5 | 168.3 ± 10.68 | 175.8 ± 35.44 | ns | 0.7302 |
| IL-6 | 4709 ± 612.7 | 4939 ± 1178 | ns | 0.7778 |
| IL-10 | 119.8 ± 10.87 | 97.63 ± 24.16 | ns | 0.3175 |
| IL-12p70 | 694.9 ± 64.09 | 680.4 ± 162.1 | ns | 0.9524 |
| IL-13 | 120.4 ± 9.005 | 110.3 ± 24.29 | ns | 0.5079 |
| IL-17A | 131.6 ± 20.37 | 121.1 ± 37.25 | ns | 0.9524 |
| IL-18 | 345 ± 49.37 | 295.2 ± 73.31 | ns | 0.5556 |
| IP-10 | 346.5 ± 17.07 | 303.7 ± 16.15 | ns | 0.0714 |
| leptin | 10505 ± 2870 | 7556 ± 3068 | ns | 0.4127 |
| LIX | 1498 ± 537.3 | 2480 ± 380.8 | ns | 0.1905 |
| MCP-1 | 1300 ± 108.8 | 1216 ± 252.8 | ns | >0.9999 |
| MIP-1α | 21.07 ± 1.806 | 20.64 ± 4.91 | ns | 0.9048 |
| MIP-2 | 174.3 ± 8.879 | 191 ± 35.32 | ns | 0.254 |
| RANTES | 1523 ± 390.4 | 2625 ± 823.6 | ns | 0.9048 |
| TNF-α | 24.14 ± 1.933 | 23.82 ± 6.386 | ns | 0.7857 |
| VEGF | 140.4 ± 21.14 | 124.6 ± 12.97 | ns | 0.9048 |

**Supplemental Figure 10:**
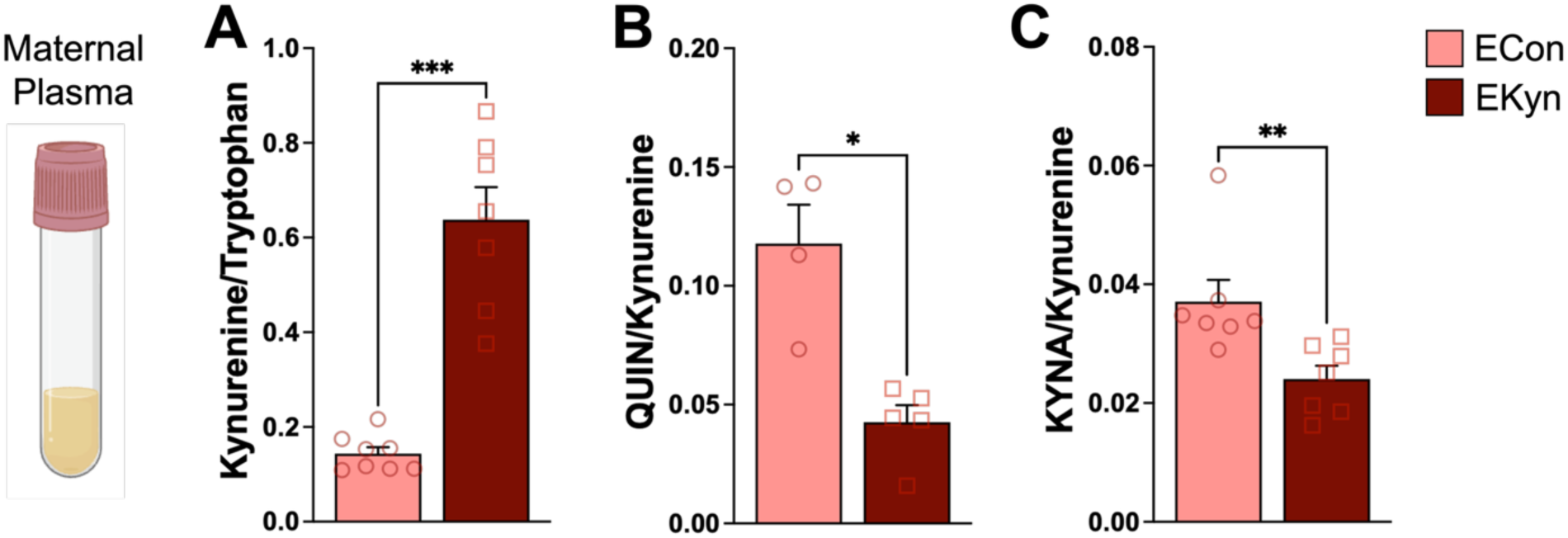
EKyn diet shifts kynurenine pathway metabolic balance in maternal plasma. Maternal plasma **(A)** Kynurenine/tryptophan ratio, **(B)** QUIN/kynurenine ratio, **(C)** KYNA/kynurenine ratio. Data are mean ± SEM with individual data points shown. Mann-Whitney test: * *p* < 0.05, ** *p* < 0.01, *** *p* < 0.001. N = 4-8 dams/group.

**Supplemental Table 3:**
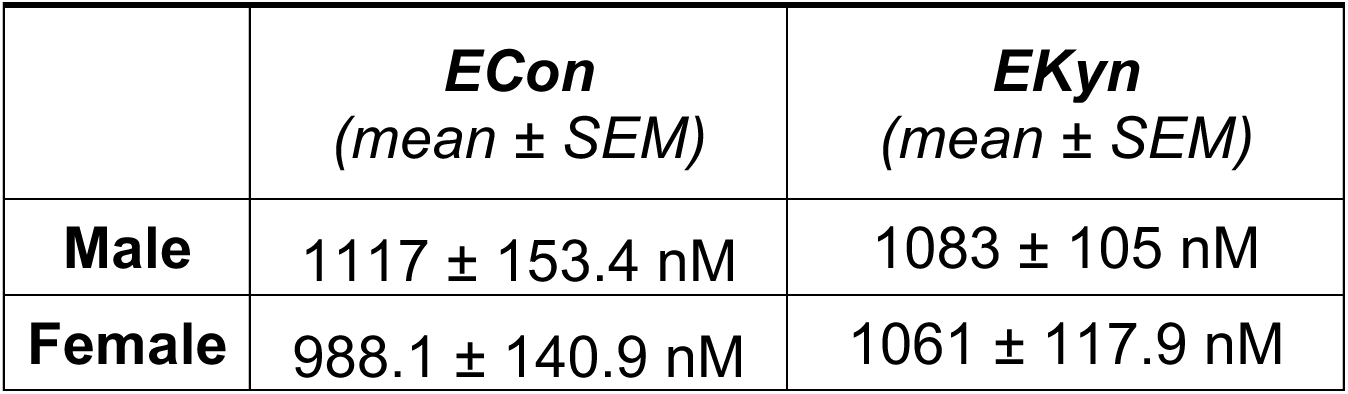
Placenta Tryptophan.

| Figure | Statistical Test | Interaction | Results | Main Effects | Results | Post-hoc Test | Post-hoc Results: |
| --- | --- | --- | --- | --- | --- | --- | --- |
| 3B | 2-way ANOVA | SleepFrag x ED | F (1, 16) = 0.8848, P = 0.3609 | ED<br>SleepFrag | F (1, 16) = 0.02566, P = 0.8748<br>F (1, 16) = 0.01396, P = 0.9074 | NA |  |
| 3D | 2-way ANOVA | SleepFrag x ED | ** F (1, 14) = 15.82, P = 0.0014 | ED<br>SleepFrag | F (1, 14) = 0.01224, P = 0.9135<br>* F (1, 14) = 6.521, P = 0.0230 | Fisher's LSD | <u>ED 18 vs ED 21:</u><br>* Control, p = 0.0206<br>** SleepFrag, p = 0.0077<br><u>Control vs SleepFrag:</u><br>*** ED 21, p = 0.0005 |
| 3E | 2-way ANOVA | SleepFrag x ED | F (1, 15) = 0.0387, P = 0.8467 | ED<br>SleepFrag | * F (1, 15) = 7.114, P = 0.0176<br>F (1, 15) = 0.2936, P = 0.5959 | Fisher's LSD |  |
| 3F | 2-way ANOVA | SleepFrag x ED | ** F (1, 16) = 10.05, P = 0.0059 | ED<br>SleepFrag | F (1, 16) = 0.05085, P = 0.8244<br>* F (1, 16) = 5.246, P = 0.0359 | Fisher's LSD | <u>ED 18 vs ED 21:</u><br>* Control, p = 0.0435<br>* SleepFrag, p = 0.0334<br><u>Control vs SleepFrag:</u><br>** ED 21, p = 0.0014 |
| 3G | 2-way ANOVA | SleepFrag x ED | ** F (1, 16) = 10.64, P = 0.0049 | ED<br>SleepFrag | F (1, 16) = 0.1400, P = 0.7132<br>F (1, 16) = 4.002, P = 0.0627 | Fisher's LSD | <u>ED 18 vs ED 21:</u><br>* Control, p = 0.0321<br>* SleepFrag, p = 0.0364<br><u>Control vs SleepFrag:</u><br>** ED 21, p = 0.0019 |
| 4A | 2-way ANOVA | SleepFrag x ED | F (1, 16) = 0.7873, P = 0.3873 | ED<br>SleepFrag | **** F (1, 17) = 26.57, P < 0.0001<br>F (1, 17) = 0.2783, P = 0.6046 | Fisher's LSD | <u>ED 18 vs ED 21:</u><br>* Control, p = 0.0147<br>*** SleepFrag, p = 0.0001 |
| 4B | 2-way ANOVA | SleepFrag x ED | F (1, 16) = 1.692, P = 0.2117 | ED<br>SleepFrag | F (1, 16) = 0.4212, P = 0.5255<br>F (1, 16) = 0.5606, P = 0.4649 | NA |  |
| 4C | 2-way ANOVA | SleepFrag x ED | ** F (1, 16) = 16.08, P = 0.0010 | ED<br>SleepFrag | *** F (1, 16) = 19.27, P = 0.0005<br>**** F (1, 16) = 29.19, P < 0.0001 | Fisher's LSD | <u>ED 18 vs ED 21:</u><br>**** Control, p < 0.0001<br><u>Control vs SleepFrag:</u><br>**** ED 21, p < 0.0001 |
| 4D | 2-way ANOVA | SleepFrag x ED | * F (1, 16) = 5.223, P = 0.0363 | ED<br>SleepFrag | F (1, 16) = 0.3776, P = 0.5475<br>F (1, 16) = 0.6196, P = 0.4427 | Fisher's LSD | <u>Control vs SleepFrag:</u><br>* ED 21, p = 0.0452 |
| 4E | Hypothalamus:<br>2-way ANOVA | SleepFrag x ED | F (1, 8) = 0.0668, P = 0.8026 | ED<br>SleepFrag | F (1, 8) = 0.2901, P = 0.6048<br>F (1, 8) = 1.119, P = 0.3210 | NA |  |
|  | Frontal Cortex:<br>2-way ANOVA | SleepFrag x ED | * F (1, 8) = 5.839, P = 0.0421 | ED<br>SleepFrag | F (1, 8) = 0.004515, P = 0.9481<br>F (1, 8) = 0.4319, P = 0.5295 | NA |  |
|  | Hippocampus:<br>2-way ANOVA | SleepFrag x ED | F (1, 15) = 3.031, P = 0.1022 | ED<br>SleepFrag | * F (1, 15) = 5.163, P = 0.0382<br>F (1, 15) = 1.707, P = 0.211 | Fisher's LSD | <u>ED 18 vs ED 21:</u><br>* Control, p = 0.0185 |
| 4F | Hypothalamus:<br>2-way ANOVA | SleepFrag x ED | F (1, 7) = 1.190, P = 0.3115 | ED<br>SleepFrag | F (1, 7) = 5.108, P = 0.0583<br>F (1, 8) = 0.1680, P = 0.6926 | NA |  |
|  | Frontal Cortex:<br>2-way ANOVA | SleepFrag x ED | F (1, 8) = 0.7404, P = 0.4146 | ED<br>SleepFrag | F (1,8) = 1.126, P = 0.3196<br>F (1, 8) = 0.2511, P = 0.6298 | NA |  |
|  | Hippocampus:<br>2-way ANOVA | SleepFrag x ED | F (1, 8) = 0.4886, P = 0.5044 | ED<br>SleepFrag | F (1, 8) = 0.8791, P = 0.5701<br>F (1, 8) = 1.550, P = 0.2483 | NA |  |
| 5A | 3-way ANOVA | SleepFrag x ED x Sex<br>SleepFrag x ED<br>ED x Sex<br>SleepFrag x Sex | F (1, 32) = 3.466, P = 0.0719<br>F (1, 32) = 2.414, P = 0.1301<br>F (1, 32) = 1.221, P = 0.2774<br>F (1, 32) = 0.09133, P = 0.7645 | ED<br>SleepFrag<br>Sex | F (1, 32) = 0.06493, P = 0.8005<br>F (1, 32) = 0.2663, P = 0.6093<br>F (1, 32) = 0.7452, P = 0.3944 | NA |  |
|  | ED 18:<br>2-way ANOVA | SleepFrag x Sex | F (1, 15) = 2.085, P = 0.1693 | SleepFrag<br>Sex | F (1, 15) = 0.9232, P = 0.3519<br>F (1, 15) = 0.05012, P = 0.8259 | NA |  |
|  | ED 21:<br>2-way ANOVA | SleepFrag x Sex | F (1, 17) = 1.808, P = 0.1964 | SleepFrag<br>Sex | F (1, 17) = 1.654, P = 0.2156<br>F (1, 17) = 1.496, P = 0.2380 | NA |  |
| 5B | 3-way ANOVA | SleepFrag x ED x Sex<br>SleepFrag x ED<br>ED x Sex<br>SleepFrag x Sex | F (1, 33) = 0.1227, P = 0.7294<br>F (1, 33) = 0.0001576, P = 0.9901<br>F (1, 33) = 0.1227, P = 0.7284<br>F (1, 33) = 0.09328, P = 0.7620 | ED<br>SleepFrag<br>Sex | F (1, 33) = 0.6938, P = 0.4109<br>F (1, 33) = 1.819, P = 0.1867<br>F (1, 33) = 0.2567, P = 0.6158 | NA |  |
|  | ED 18:<br>2-way ANOVA | SleepFrag x Sex | F (1, 16) = 0.789, P = 0.3875 | SleepFrag<br>Sex | F (1, 16) = 0.750, P = 0.3992<br>F (1, 16) = 0.308, P = 0.5862 | NA |  |
|  | ED 21:<br>2-way ANOVA | SleepFrag x Sex | F (1, 17) = 2.362, P = 0.1427 | SleepFrag<br>Sex | F (1, 17) = 1.115, P = 0.3058<br>F (1, 17) = 0.014, P = 0.9049 | NA |  |
| 5C | 3-way ANOVA | SleepFrag x ED x Sex<br>SleepFrag x ED<br>ED x Sex<br>SleepFrag x Sex | F (1, 33) = 3.855, P = 0.0581<br>** F (1, 33) = 8.820, P = 0.0055<br>F (1, 33) = 2.878, P = 0.0992<br>F (1, 33) = 3.036, P = 0.0907 | ED<br>SleepFrag<br>Sex | F (1, 33) = 0.3191, P = 0.5760<br>F (1, 33) = 1.056, P = 0.3115<br>F (1, 33) = 0.1789, P = 0.6751 | NA |  |
|  | ED 18:<br>2-way ANOVA | SleepFrag x Sex | F (1, 16) = 0.03604, P = 0.8518 | SleepFrag<br>Sex | F (1, 16) = 2.781, P = 0.1148<br>F (1, 16) = 1.196, P = 0.2904 | NA |  |
|  | ED 21:<br>2-way ANOVA | SleepFrag x Sex | * F (1, 17) = 5.423, P = 0.0325 | SleepFrag<br>Sex | * F (1, 17) = 6.310, P = 0.0224<br>F (1, 17) = 1.773, P = 0.2005 | Fisher's LSD | <u>Control vs SleepFrag:</u><br>** Male, p = 0.0041<br><u>Male vs Female:</u><br>* SleepFrag, p = 0.0122 |
| 5D | 3-way ANOVA | SleepFrag x ED x Sex<br>SleepFrag x ED<br>ED x Sex<br>SleepFrag x Sex | F (1, 50) = 0.1321, P = 0.7178<br>F (1, 50) = 2.719, P = 0.1054<br>F (1, 50) = 0.3215, P = 0.5733<br>F (1, 50) = 0.0009230, P = 0.9759 | ED<br>SleepFrag<br>Sex | F (1, 50) = 0.04280, P = 0.8369<br>F (1, 50) = 3.534, P = 0.0659<br>F (1, 50) = 1.470, P = 0.2310 | NA |  |
|  | ED 18:<br>2-way ANOVA | SleepFrag x Sex | F (1, 20) = 0.0808, P = 0.7791 | SleepFrag<br>Sex | F (1, 20) = 0.03884, P = 0.8457<br>F (1, 20) = 2.306, P = 0.1445 | NA |  |
|  | ED 21:<br>2-way ANOVA | SleepFrag x Sex | F (1, 30) = 0.07759, P = 0.7825 | SleepFrag<br>Sex | * F (1, 30) = 6.229, P = 0.0183<br>F (1, 30) = 0.2085, P = 0.6513 | NA |  |
| 5E | 3-way ANOVA | SleepFrag x ED x Sex<br>SleepFrag x ED<br>ED x Sex<br>SleepFrag x Sex | F (1, 50) = 0.1013, P = 0.7516<br>F (1, 50) = 0.4506, P = 0.5051<br>F (1, 50) = 0.03365, P = 0.8552<br>F (1, 50) = 0.2944, P = 0.5898 | ED<br>SleepFrag<br>Sex | F (1, 50) = 1.905, P = 0.1737<br>**** F (1, 50) = 34.36, P < 0.0001<br>F (1, 50) = 0.2539, P = 0.6166 | NA |  |
|  | ED 18:<br>2-way ANOVA | SleepFrag x Sex | F (1, 20) = 0.02023, P = 0.8883 | SleepFrag<br>Sex | ** F (1, 20) = 10.83, P = 0.0037<br>F (1, 20) = 0.04126, P = 0.8411 | Fisher's LSD | <u>Control vs SleepFrag:</u><br>* male, p = 0.0248<br>* female, p = 0.0377 |
|  | ED 21:<br>2-way ANOVA | SleepFrag x Sex | F (1, 30) = 0.4839, P = 0.4920 | SleepFrag<br>Sex | **** F (1, 30) = 27.87, P < 0.0001<br>F (1, 30) = 0.3085, P = 0.5827 | Fisher's LSD | <u>Control vs SleepFrag:</u><br>*** male, p = 0.0002<br>** female, p = 0.0029 |
| 5F | 3-way ANOVA | SleepFrag x ED x Sex<br>SleepFrag x ED<br>ED x Sex<br>SleepFrag x Sex | * F (1, 50) = 4.861, P = 0.0321<br>F (1, 50) = 0.08130, P = 0.7767<br>* F (1, 50) = 5.840, P = 0.0194<br>F (1, 50) = 1.490, P = 0.2279 | ED<br>SleepFrag<br>Sex | **** F (1, 50) = 65.68, P < 0.0001<br>F (1, 50) = 2.150, P = 0.1488<br>** F (1, 50) = 8.623, P = 0.0050 | NA |  |
|  | ED 18:<br>2-way ANOVA | SleepFrag x Sex | F (1, 20) = 3.108, P = 0.0932 | SleepFrag<br>Sex | F (1, 20) = 0.3696, P = 0.5501<br>* F (1, 20) = 7591, P = 0.0122 | Fisher's LSD | <u>Male vs Female:</u><br>* Control, p = 0.0119 |
|  | ED 21:<br>2-way ANOVA | SleepFrag x Sex | F (1, 30) = 0.9765, P = 0.3310 | SleepFrag<br>Sex | F (1, 30) = 3.095, P = 0.0887<br>F (1, 30) = 0.2725, P = 0.6055 | NA |  |
| 6A | 3-way ANOVA | SleepFrag x ED x Sex<br>SleepFrag x ED<br>ED x Sex<br>SleepFrag x Sex | F (1, 61) = 0.01939, P = 0.8897<br>** F (1, 61) = 8.911, P = 0.0041<br>F (1, 61) = 0.001944, P = 0.9650<br>F (1, 61) = 0.05127, P = 0.8216 | ED<br>SleepFrag<br>Sex | **** F (1, 61) = 36.53, P < 0.0001<br>*** F (1, 61) = 12.54, P = 0.0008<br>F (1, 61) = 0.02879, P = 0.8658 | NA |  |
|  | ED 18:<br>2-way ANOVA | SleepFrag x Sex | F (1, 31) = 0.005402, P = 0.9419 | SleepFrag<br>Sex | F (1, 31) = 0.22, P = 0.6423<br>F (1, 31) = 0.01121, P = 0.9164 | NA |  |
|  | ED 21:<br>2-way ANOVA | SleepFrag x Sex | F (1, 30) = 0.05302, P = 0.8194 | SleepFrag<br>Sex | *** F (1, 30) = 16.89, P = 0.0003<br>F (1, 30) = 0.01812, P = 0.8938 | Fisher's LSD | <u>Control vs SleepFrag:</u><br>** male, p = 0.0045<br>* female, p = 0.0102 |
| 6B | 3-way ANOVA | SleepFrag x ED x Sex<br>SleepFrag x ED<br>ED x Sex<br>SleepFrag x Sex | F (1, 60) = 1.142, P = 0.2896<br>F (1, 60) = 1.796, P = 0.1852<br>F (1, 60) = 0.1512, P = 0.6988<br>F (1, 60) = 0.2221, P = 0.6391 | ED<br>SleepFrag<br>Sex | F (1, 60) = 1.328, P = 0.2538<br>** F (1, 60) = 9.812, P = 0.0027<br>F (1, 60) = 0.1856, P = 0.6682 | NA |  |
|  | ED 18:<br>2-way ANOVA | SleepFrag x Sex | F (1, 31) = 0.8678, P = 0.3588 | SleepFrag<br>Sex | F (1, 31) = 1.176, P = 0.2866<br>F (1, 31) = 0.2459, P = 0.6235 | NA |  |
|  | ED 21:<br>2-way ANOVA | SleepFrag x Sex | F (1, 29) = 0.2768, P = 0.6028 | SleepFrag<br>Sex | *** F (1, 29) = 15.53, P = 0.0005<br>F (1, 29) = 0.001368, P = 0.9707 | Fisher's LSD | <u>Control vs SleepFrag:</u><br>* male, p = 0.0246<br>** female, p = 0.0032 |
| 6C | 3-way ANOVA | SleepFrag x ED x Sex<br>SleepFrag x ED<br>ED x Sex<br>SleepFrag x Sex | F (1, 58) = 0.0877, P = 0.7681<br>* F (1, 58) = 4.242, P = 0.0439<br>F (1, 58) = 0.9468, P = 0.3346<br>F (1, 58) = 0.4406, P = 0.5095 | ED<br>SleepFrag<br>Sex | **** F (1, 58) = 31.82, P < 0.0001<br>F (1, 58) = 3.959, P = 0.0513<br>F (1, 58) = 1.289, P = 0.2609 | NA |  |
|  | ED 18:<br>2-way ANOVA | SleepFrag x Sex | F (1, 30) = 0.1730, P = 0.6804 | SleepFrag<br>Sex | F (1, 30) = 0.006255, P = 0.9375<br>F (1, 30) = 0.03378, P = 0.8554 | NA |  |
|  | ED 21:<br>2-way ANOVA | SleepFrag x Sex | F (1, 28) = 0.2862, P = 0.5969 | SleepFrag<br>Sex | * F (1, 28) = 5.091, P = 0.0320<br>F (1, 28) = 1.380, P = 0.2499 | Fisher's LSD | <u>Control vs SleepFrag:</u><br>male, p = 0.0680 |
| 7B | 3-way RM ANOVA | EKyn x vigilant state x light phase<br>EKyn x vigilant state<br>EKyn x light phase<br>vigilant state x light phase | ** F (1, 18) = 14.58, P = 0.0013<br>F (1, 20) = 0.9880, P = 0.3321<br>F (1, 18) = 0.000004965, P = 0.9982<br>**** F (1, 18) = 1025, P < 0.0001 | EKyn<br>vigilant state<br>light phase | F (1, 20) = 0.000009, P = 0.9976<br>**** F (1, 20) = 61.67, P < 0.0001<br>F (1, 18) = 0.000004965, P = 0.9982 | NA |  |
|  | Light Phase:<br>2-way RM ANOVA | EKyn x vigilant state | F (1, 10) = 0.74966, P = 0.4069 | EKyn<br>vigilant state | * F (1, 10) = 5.00, P = 0.0493<br>**** F (1, 10) = 260.7, P < 0.0001 | Fisher's LSD |  |
|  | Dark Phase:<br>2-way RM ANOVA | EKyn x vigilant state | * F (1, 9) = 5.5, P = 0.0436 | EKyn<br>vigilant state | F (1, 9) = 0.8182, P = 0.3893<br>**** F (1, 9) = 77.67, P < 0.0001 | Fisher's LSD | <u>ECon vs EKyn:</u><br>* wake, p = 0.0306<br>* sleep, p = 0.0307 |
| 7C | 2-way RM ANOVA | EKyn x light phase | F (1, 8) = 2.139, P = 0.1818 | EKyn<br>light phase | * F (1, 9) = 6.128, P = 0.0352<br>**** F (1, 8) = 66.07, P < 0.0001 | Fisher's LSD | <u>ECon vs EKyn:</u><br>* dark phase, p = 0.01 |
| 9H | 2-way ANOVA | EKyn x Sex | F (1, 31) = 0.09940, P = 0.7547 | EKyn<br>Sex | **** F (1, 31) = 110.0, P < 0.0001<br>F (1, 31) = 0.2552, P = 0.6170 | Fisher's LSD | <u>ECon vs EKyn:</u><br>**** male, p < 0.0001<br>**** female, p < 0.0001 |
| 9I | 2-way ANOVA | EKyn x Sex | F (1, 29) = 0.2095, P = 0.6506 | EKyn<br>Sex | **** F (1, 29) = 80.50, P < 0.0001<br>* F (1, 29) = 5.274, P = 0.0291 | Fisher's LSD | <u>ECon vs EKyn:</u><br>**** male, p < 0.0001<br>**** female, p < 0.0001 |
| 9J | 2-way ANOVA | EKyn x Sex | F (1, 29) = 0.005448, P = 0.9417 | EKyn<br>Sex | F (1, 29) = 0.04081, P = 0.8413<br>F (1, 29) = 1.096, P = 0.3038 | NA |  |
| 9K | 2-way ANOVA | EKyn x Sex | * F (1, 34) = 5.647, P = 0.0233 | EKyn<br>Sex | **** F (1, 34) = 228.1, P < 0.0001<br>* F (1, 34) = 6.666, P = 0.0143 | Fisher's LSD | <u>ECon vs EKyn:</u><br>**** male, p < 0.0001<br>**** female, p < 0.0001<br><u>Male vs Female:</u><br>** EKyn, p < 0.01 |
| 9L | 2-way ANOVA | EKyn x Sex | F (1, 32) = 0.2158, P = 0.6454 | EKyn<br>Sex | **** F (1, 32) = 101.8, P < 0.0001<br>F (1, 32) = 0.3358, P = 0.5663 | Fisher's LSD | <u>ECon vs EKyn:</u><br>**** male, p < 0.0001<br>**** female, p < 0.0001 |
| 9M | 2-way ANOVA | EKyn x Sex | F (1, 29) = 1.008, P = 0.3238 | EKyn<br>Sex | F (1, 29) = 0.2314, p = 0.6341<br>* F (1, 29) = 5.104, p = 0.0316 | Fisher's LSD | <u>ECon vs EKyn:</u><br>* male, p = 0.0187 |

| Supplemental Figure | Statistical Test | Interaction | Results | Main Effects | Results | Post-hoc Test | Post-hoc Results: |
| --- | --- | --- | --- | --- | --- | --- | --- |
| S3A | 2-way ANOVA | SleepFrag x ED | ** F (1, 16) = 10.11, p = 0.0058 | ED<br>SleepFrag | F (1, 16) = 0.2264, p = 0.6406<br>* F (1, 16) = 6.706, p = 0.0198 | Fisher's LSD | <u>ED 18 vs ED 21:</u><br>* SleepFrag, p = 0.0107<br><u>Control vs SleepFrag:</u><br>*** ED 21, p = 0.0009 |
| S3B | 2-way ANOVA | SleepFrag x ED | ** F (1, 16) = 8.838, p = 0.0090 | ED<br>SleepFrag | F (1, 16) = 0.1101, p = 0.7443<br>* F (1, 16) = 8.108, p = 0.0116 | Fisher's LSD | <u>ED 18 vs ED 21:</u><br>* SleepFrag, p = 0.0189<br><u>Control vs SleepFrag:</u><br>*** ED 21, p = 0.0008 |
| S3C | 2-way ANOVA | SleepFrag x ED | * F (1, 16) = 7.257, p = 0.0160 | ED<br>SleepFrag | F (1, 16) = 0.0009397, p = 0.9759<br>F (1, 16) = 2.327, p = 0.1467 | Fisher's LSD | <u>Control vs SleepFrag:</u><br>** ED 21, p = 0.0088 |
| S3D | 2-way ANOVA | SleepFrag x ED | * F (1, 14) = 7.424, p = 0.0164 | ED<br>SleepFrag | F (1, 14) = 0.004908, p = 0.9451<br>F (1, 14) = 3.698, p = 0.0751 | Fisher's LSD | <u>Control vs SleepFrag:</u><br>** ED 21, p = 0.0064 |
| S3E | 2-way ANOVA | SleepFrag x ED | * F (1, 16) = 6.892, p = 0.0184 | ED<br>SleepFrag | F (1, 16) = 0.2787, p = 0.6048<br>F (1, 16) = 1.929, p = 0.1839 | Fisher's LSD | <u>ED 18 vs ED 21:</u><br>* SleepFrag, p = 0.0240<br><u>Control vs SleepFrag:</u><br>* ED 21, p = 0.0119 |
| S3F | 2-way ANOVA | SleepFrag x ED | * F (1, 16) = 7.684, p = 0.0136 | ED<br>SleepFrag | F (1, 16) = 0.0005177, p = 0.9821<br>F (1, 16) = 3.859, p = 0.0671 | Fisher's LSD | <u>ED 18 vs ED 21:</u><br>* SleepFrag, p = 0.0451<br><u>Control vs SleepFrag:</u><br>** ED 21, p = 0.0041 |
| S3G | 2-way ANOVA | SleepFrag x ED | ** F (1, 16) = 10.79, p = 0.0047 | ED<br>SleepFrag | F (1, 16) = 0.3116, p = 0.5844<br>* F (1, 16) = 5.834, p = 0.0280 | Fisher's LSD | <u>ED 18 vs ED 21:</u><br>* Control, p = 0.0246<br>* SleepFrag, p = 0.0467<br><u>Control vs SleepFrag:</u><br>*** ED 21, p = 0.0010 |
| S3I | 2-way ANOVA | SleepFrag x ED | * F (1, 16) = 7.578, p = 0.0142 | ED<br>SleepFrag | F (1, 16) = 0.004338, p = 0.9483<br>F (1, 16) = 3.0, p = 0.1025 | Fisher's LSD | <u>ED 18 vs ED 21:</u><br>* SleepFrag, p = 0.0496<br><u>Control vs SleepFrag:</u><br>** ED 21, p = 0.0059 |
| S3J | 2-way ANOVA | SleepFrag x ED | * F (1, 16) = 7.250, p = 0.0160 | ED<br>SleepFrag | F (1, 16) = 0.03483, p = 0.8543<br>* F (1, 16) = 4.710, p = 0.0454 | Fisher's LSD | <u>ED 18 vs ED 21:</u><br>* SleepFrag, p = 0.0369<br><u>Control vs SleepFrag:</u><br>** ED 21, p = 0.0034 |
| S3K | 2-way ANOVA | SleepFrag x ED | ** F (1, 16) = 13.57, p = 0.0020 | ED<br>SleepFrag | F (1, 16) = 0.4068, p = 0.5324<br>F (1, 16) = 2.206, p = 0.1569 | Fisher's LSD | <u>ED 18 vs ED 21:</u><br>** SleepFrag, p = 0.0035<br><u>Control vs SleepFrag:</u><br>** ED 21, p = 0.0021 |
| S3L | 2-way ANOVA | SleepFrag x ED | * F (1, 16) = 6.263, p = 0.0235 | ED<br>SleepFrag | F (1, 16) = 0.01679, p = 0.8985<br>F (1, 16) = 1.212, p = 0.2873 | Fisher's LSD | <u>Control vs SleepFrag:</u><br>* ED 21, p = 0.0215 |
| S3M | 2-way ANOVA | SleepFrag x ED | F (1, 15) = 0.1230, p = 0.7307 | ED<br>SleepFrag | * F (1, 15) = 7.857, p = 0.0134<br>*** F (1, 15) = 18.74, p = 0.0006 | Fisher's LSD | <u>Control vs SleepFrag:</u><br>** ED 18, p = 0.0054<br>* ED 21, p = 0.0117 |
| S3N | 2-way ANOVA | SleepFrag x ED | ** F (1, 16) = 9.148, p = 0.0081 | ED<br>SleepFrag | * F (1, 16) = 8.122, p = 0.0116<br>F (1, 16) = 0.5617, p = 0.4645 | Fisher's LSD | <u>ED 18 vs ED 21:</u><br>*** SleepFrag, p = 0.0007<br><u>Control vs SleepFrag:</u><br>** ED 21, p = 0.0088 |
| S3O | 2-way ANOVA | SleepFrag x ED | ** F (1, 16) = 10.32, p = 0.0054 | ED<br>SleepFrag | F (1, 16) = 0.03310, p = 0.8579<br>F (1, 16) = 3.551, p = 0.0778 | Fisher's LSD | <u>ED 18 vs ED 21:</u><br>* Control, p = 0.0436<br>* SleepFrag, p = 0.0292<br><u>Control vs SleepFrag:</u><br>** ED 21, p = 0.0024 |
| S4A | 2-way ANOVA | SleepFrag x ED | F (1, 16) = 0.03544, p = 0.8531 | ED<br>SleepFrag | F (1, 16) = 1.840, p = 0.1938<br>F (1, 16) = 0.4210, p = 0.5256 | NA |  |
| S4B | 2-way ANOVA | SleepFrag x ED | F (1, 16) = 2.292, p = 0.1496 | ED<br>SleepFrag | F (1, 16) = 0.7981, p = 0.3849<br>F (1, 16) = 0.01918, p = 0.8916 | NA |  |
| S4C | 2-way ANOVA | SleepFrag x ED | * F (1, 16) = 4.961, p = 0.0406 | ED<br>SleepFrag | ** F (1, 16) = 9.147, p = 0.0081<br>* F (1, 16) = 8.105, p = 0.0117 | Fisher's LSD | <u>ED 18 vs ED 21:</u><br>** Control, p = 0.0037<br><u>Control vs SleepFrag:</u><br>** ED 21, p = 0.0025 |
| S4D | 2-way ANOVA | SleepFrag x ED | F (1, 16) = 0.04998, p = 0.08259 | ED<br>SleepFrag | F (1, 16) = 3.918, p = 0.0652<br>F (1, 16) = 4.320, p = 0.0541 | NA |  |
| S5A | 3-way ANOVA | SleepFrag x ED x Sex<br>SleepFrag x ED<br>ED x Sex<br>SleepFrag x Sex | F (1, 32) = 0.004903, p = 0.9556<br>F (1, 32) = 1.083, p = 0.3508<br>F (1, 32) = 3.053, p = 0.0902<br>F (1, 32) = 0.1593, p = 0.6924 | ED<br>SleepFrag<br>Sex | **** F (1, 32) = 1941, p < 0.0001<br>F (1, 32) = 3.026, p = 0.0915<br>** F (1, 32) = 0.810, p = 0.0037 | NA |  |
|  | ED 18:<br>2-way ANOVA | SleepFrag x Sex | F (1, 16) = 0.044, p = 0.8365 | SleepFrag<br>Sex | F (1, 16) = 0.1984, p = 0.6620<br>F (1, 16) = 0.3905, p = 0.7790 | NA |  |
|  | ED 21:<br>2-way ANOVA | SleepFrag x Sex | F (1, 16) = 0.1431, p = 0.7102 | SleepFrag<br>Sex | * F (1, 16) = 5.025, p = 0.0395<br>** F (1, 16) = 15.48, p = 0.0012 | Fisher's LSD |  |
| S5B | 3-way ANOVA | SleepFrag x ED x Sex<br>SleepFrag x ED<br>ED x Sex<br>SleepFrag x Sex | F (1, 32) = 0.1161, p = 0.7356<br>F (1, 32) = 0.3393, p = 0.5643<br>F (1, 32) = 0.1488, p = 0.7022<br>F (1, 32) = 0.03246, p = 0.8581 | ED<br>SleepFrag<br>Sex | **** F (1, 32) = 48.84, p < 0.0001<br>F (1, 32) = 1.472, p = 0.2338<br>F (1, 32) = 0.7980, p = 0.2338 | NA |  |
|  | ED 18:<br>2-way ANOVA | SleepFrag x Sex | F (1, 16) = 0.1074, p = 0.7474 | SleepFrag<br>Sex | F (1, 16) = 0.1576, p = 0.6966<br>F (1, 16) = 0.1020, p = 0.7536 | NA |  |
|  | ED 21:<br>2-way ANOVA | SleepFrag x Sex | F (1, 16) = 2.189, p = 0.1585 | SleepFrag<br>Sex | F (1, 16) = 0.01748, p = 0.8945<br>F (1, 16) = 1.110, p = 0.3077 | NA |  |
| S5C | 3-way ANOVA | SleepFrag x ED x Sex<br>SleepFrag x ED<br>ED x Sex<br>SleepFrag x Sex | F (1, 32) = 0.0503, p = 0.8240<br>F (1, 32) = 0.03202, p = 0.8591<br>F (1, 32) = 0.01019, p = 0.9209<br>F (1, 32) = 0.04209, p = 0.8387 | ED<br>SleepFrag<br>Sex | **** F (1, 32) = 391.6, p < 0.0001<br>F (1, 32) = 0.1627, p = 0.6894<br>F (1, 32) = 0.6215, p = 0.4363 | NA |  |
|  | ED 18:<br>2-way ANOVA | SleepFrag x Sex | F (1, 16) = 0.1656, p = 0.6894 | SleepFrag<br>Sex | F (1, 16) = 0.04521, p = 0.8343<br>F (1, 16) = 0.4244, p = 0.5240 | NA |  |
|  | ED 21:<br>2-way ANOVA | SleepFrag x Sex | F (1, 16) = 0.0001265, p = 0.9912 | SleepFrag<br>Sex | F (1, 16) = 0.1174, p = 0.7363<br>F (1, 16) = 0.2740, p = 0.6079 | NA |  |
| S5D | 2-way ANOVA | ED x SleepFrag | F (1, 16) = 0.1325, p = 0.7206 | ED<br>SleepFrag | F (1, 16) = 0.2830, p = 0.6021<br>F (1, 16) = 0.2830, p = 0.6021 | NA |  |
| S6A | 3-way ANOVA | SleepFrag x ED x Sex<br>SleepFrag x ED<br>ED x Sex<br>SleepFrag x Sex | F (1, 54) = 0.2056, p = 0.6520<br>F (1, 54) = 2.122, p = 0.1510<br>F (1, 54) = 0.4052, p = 0.5271<br>F (1, 54) = 0.03257, p = 0.8575 | ED<br>SleepFrag<br>Sex | **** F (1, 54) = 66.98, p < 0.0001<br>F (1, 54) = 1.511, p = 0.2254<br>F (1, 54) = 0.000157, p = 0.990 | NA |  |
|  | ED 18:<br>2-way ANOVA | SleepFrag x Sex | F (1, 24) = 0.05513, p = 0.8164 | SleepFrag<br>Sex | F (1, 24) = 0.03831, p = 0.8465<br>F (1, 24) = 0.3117, p = 0.5818 | NA |  |
|  | ED 21:<br>2-way ANOVA | SleepFrag x Sex | F (1, 30) = 0.1778, p = 0.6763 | SleepFrag<br>Sex | F (1, 30) = 3.192, p = 0.0841<br>F (1, 30) = 0.1722, p = 0.6811 | NA |  |
| S6B | 3-way ANOVA | SleepFrag x ED x Sex<br>SleepFrag x ED<br>ED x Sex<br>SleepFrag x Sex | * F (1, 53) = 4.562, p = 0.0373<br>F (1, 53) = 0.4585, p = 0.5013<br>F (1, 53) = 0.08293, p = 0.7745<br>F (1, 53) = 2.414, p = 0.1262 | ED<br>SleepFrag<br>Sex | **** F (1, 53) = 169.0, p < 0.0001<br>F (1, 53) = 0.002816, p = 0.9579<br>F (1, 53) = 0.5871, p = 0.4469 | NA |  |
|  | ED 18:<br>2-way ANOVA | SleepFrag x Sex | F (1, 24) = 0.944, p = 0.3409 | SleepFrag<br>Sex | F (1, 24) = 1.485, p = 0.2349<br>F (1, 24) = 0.637, p = 0.4326 | NA |  |
|  | ED 21:<br>2-way ANOVA | SleepFrag x Sex | * F (1, 29) = 4.621, p = 0.0401 | SleepFrag<br>Sex | F (1, 29) = 0.1322, 0.7188<br>F (1, 29) = 0.3773, p = 0.5438 | Fisher's LSD |  |
| S7A | 3-way ANOVA | SleepFrag x ED x Sex<br>SleepFrag x ED<br>ED x Sex<br>SleepFrag x Sex | F (1, 50) = 0.05997, p = 0.8075<br>F (1, 50) = 0.03988, p = 0.8425<br>F (1, 50) = 0.02886, p = 0.8625<br>F (1, 50) = 0.7293, p = 0.3972 | ED<br>SleepFrag<br>Sex | F (1, 50) = 0.7121, p = 0.4028<br>**** F (1, 50) = 32.99, p < 0.0001<br>F (1, 50) = 0.8844, p = 0.3515 | NA |  |
|  | ED 18:<br>2-way ANOVA | SleepFrag x Sex | F (1, 20) = 0.1210, p = 0.7315 | SleepFrag<br>Sex | ** F (1, 20) = 10.03, p = 0.0049<br>F (1, 20) = 0.4022, p = 0.5331 | Fisher's LSD | <u>Control vs SleepFrag:</u><br>* male, p = 0.0219<br>female, p = 0.0601 |
|  | ED 21:<br>2-way ANOVA | SleepFrag x Sex | F (1, 30) = 0.9367, p = 0.3409 | SleepFrag<br>Sex | **** F (1, 30) = 27.40, p < 0.0001<br>F (1, 30) = 0.4605, p = 0.5026 | Fisher's LSD | <u>Control vs SleepFrag:</u><br>*** male, p = 0.0001<br>** female, p = 0.0052 |
| S7B | 3-way ANOVA | SleepFrag x ED x Sex<br>SleepFrag x ED<br>ED x Sex<br>SleepFrag x Sex | F (1, 50) = 2.492, p = 0.1207<br>F (1, 50) = 0.6394, p = 0.4277<br>F (1, 50) = 3.315, p = 0.0746<br>F (1, 50) = 2.290, p = 0.1365 | ED<br>SleepFrag<br>Sex | **** F (1, 50) = 31.55, p < 0.0001<br>* F (1, 50) = 6.446, p = 0.0143<br>* F (1, 50) = 5.211, p = 0.0267 | NA |  |
|  | ED 18:<br>2-way ANOVA | SleepFrag x Sex | F (1, 20) = 1.881, p = 0.1854 | SleepFrag<br>Sex | F (1, 20) = 2.193, p = 0.1542<br>F (1, 20) = 3.313, p = 0.0837 | NA |  |
|  | ED 21:<br>2-way ANOVA | SleepFrag x Sex | F (1, 30) = 0.009647, p = 0.9244 | SleepFrag<br>Sex | * F (1, 30) = 6.835, p = 0.0138<br>F (1, 30) = 0.4820, p = 0.4929 | Fisher's LSD |  |
| S7C | 3-way ANOVA | SleepFrag x ED x Sex<br>SleepFrag x ED<br>ED x Sex<br>SleepFrag x Sex | F (1, 60) = 1.076, p = 0.3037<br>F (1, 60) = 0.2328, p = 0.6312<br>F (1, 60) = 0.3333, p = 0.5659<br>F (1, 60) = 0.1015, p = 0.7511 | ED<br>SleepFrag<br>Sex | **** F (1, 60) = 24.79, p < 0.0001<br>F (1, 60) = 0.01664, p = 0.8978<br>F (1, 60) = 0.06147, p = 0.8050 | NA |  |
|  | ED 18:<br>2-way ANOVA | SleepFrag x Sex | F (1, 31) = 0.5768, p = 0.4533 | SleepFrag<br>Sex | F (1, 31) = 0.1172, p = 0.7344<br>F (1, 31) = 0.2136, p = 0.6472 | NA |  |
|  | ED 21:<br>2-way ANOVA | SleepFrag x Sex | F (1, 29) = 0.6110, p = 0.4407 | SleepFrag<br>Sex | F (1, 29) = 0.1477, p = 0.7035<br>F (1, 29) = 0.1283, p = 0.7228 | NA |  |
| S7D | 3-way ANOVA | SleepFrag x ED x Sex<br>SleepFrag x ED<br>ED x Sex<br>SleepFrag x Sex | F (1, 57) = 1.318, p = 0.2558<br>F (1, 57) = 0.1497, p = 0.7003<br>F (1, 57) = 3.025, p = 0.0874<br>F (1, 57) = 2.062, p = 0.1565 | ED<br>SleepFrag<br>Sex | **** F (1, 57) = 64.34, p < 0.0001<br>F (1, 57) = 0.8992, p = 0.3470<br>F (1, 57) = 1.783, p = 0.1871 | NA |  |
|  | ED 18:<br>2-way ANOVA | SleepFrag x Sex | F (1, 29) = 0.1370, p = 0.7139 | SleepFrag<br>Sex | F (1, 29) = 2.944, p = 0.0969<br>F (1, 29) = 0.2695, p = 0.6076 | NA |  |
|  | ED 21:<br>2-way ANOVA | SleepFrag x Sex | F (1, 28) = 2.012, p = 0.1671 | SleepFrag<br>Sex | F (1, 28) = 0.09497, p = 0.7602<br>F (1, 28) = 2.849, p = 0.1026 | NA |  |
| S8A | 2-way RM ANOVA | EKyn x Light Phase | * F (1, 9) = 7.287, p = 0.0244 | EKyn<br>Light Phase | F (1, 10) = 0.497, p = 0.4968<br>**** F (1, 9) = 513.1, p < 0.0001 | Fisher's LSD |  |
| S8B | 2-way RM ANOVA | EKyn x Light Phase | * F (1, 9) = 8.944, p = 0.0152 | EKyn<br>Light Phase | F (1, 10) = 0.248, p = 0.6290<br>**** F (1, 9) = 514.0, p < 0.0001 | Fisher's LSD |  |
| S8C | 2-way RM ANOVA | EKyn x Light Phase | F (1, 8) = 0.1436, p = 0.7146 | EKyn<br>Light Phase | F (1, 10) = 0.651, p = 0.4382<br>*** F (1, 8) = 26.10, p = 0.0009 | NA |  |
| S8D | 2-way RM ANOVA | EKyn x Light Phase | F (1, 9) = 0.0119, p = 0.9152 | EKyn<br>Light Phase | F (1, 10) = 3.439, p = 0.0934<br>* F (1, 9) = 6.362, p = 0.0327 | NA |  |
| S8E | 2-way RM ANOVA | EKyn x Light Phase | F (1, 9) = 0.0953, p = 0.7646 | EKyn<br>Light Phase | F (1, 10) = 2.526, p = 0.1431<br>*** F (1, 9) = 25.39, p = 0.0007 | NA |  |
| S8F | 2-way RM ANOVA | EKyn x Light Phase | F (1, 9) = 0.0495, p = 0.8288 | EKyn<br>Light Phase | F (1, 10) = 0.967, p = 0.3484<br>**** F (1, 9) = 55.87, p < 0.0001 | NA |  |
| S8G | 2-way RM ANOVA | EKyn x Light Phase | F (1, 10) = 0.9694, p = 0.3480 | EKyn<br>Light Phase | F (1, 10) = 1.814, p = 0.2077<br>** F (1, 10) = 19.86, p = 0.0012 | NA |  |
| S8H | 2-way RM ANOVA | EKyn x Light Phase | F (1, 10) = 0.6267, p = 0.4469 | EKyn<br>Light Phase | F (1, 10) = 0.7117, p = 0.4186<br>** F (1, 10) = 13.88, p = 0.0039 | NA |  |
| S8I | 2-way RM ANOVA | EKyn x Light Phase | F (1, 10) = 0.6553, p = 0.4371 | EKyn<br>Light Phase | F (1, 10) = 0.5513, p = 0.4749<br>F (1, 10) = 0.3625, p = 0.5605 | NA |  |
| S9A | 2-way RM ANOVA | EKyn x Sex | $F(1, 26) = 0.02759, p = 0.8696$ | EKyn<br>Sex | $F(1, 26) = 3.611, p = 0.0685$<br>$F(1, 26) = 1.624, p = 0.2138$ | NA | |
| S9B | 2-way RM ANOVA | EKyn x Sex | $F(1, 26) = 0.07199, p = 0.7906$ | EKyn<br>Sex | $F(1, 26) = 0.04287, p = 0.8376$<br>$F(1, 26) = 0.1241, p = 0.7275$ | NA | |
| S9C | 2-way RM ANOVA | EKyn x Sex | $F(1, 26) = 0.07350, p = 0.7884$ | EKyn<br>Sex | $F(1, 26) = 0.5813, p = 0.4527$<br>$F(1, 26) = 0.01667, p = 0.8983$ | NA | |
| Supplemental Table 3 | 2-way RM ANOVA | EKyn x Sex | $F(1, 24) = 0.1478, p = 0.7041$ | EKyn<br>Sex | $F(1, 24) = 0.02014, p = 0.8883$<br>$F(1, 24) = 0.2937, p = 0.5929$ | NA | |

